# HIV-1 budding requires cortical actin disassembly by the oxidoreductase MICAL1

**DOI:** 10.1101/2024.10.07.616958

**Authors:** Thomas Serrano, Nicoletta Casartelli, Foad Ghasemi, Hugo Wioland, Frédérique Cuvelier, Audrey Salles, Maryse Moya-Nilges, Lisa Welker, Serena Bernacchi, Marc Ruff, Antoine Jégou, Guillaume Romet-Lemonne, Olivier Schwartz, Stéphane Frémont, Arnaud Echard

## Abstract

Many enveloped viruses bud from the plasma membrane that is tightly associated with a dense and thick actin cortex. This actin network represents a significant challenge for membrane deformation and scission, and how it is remodeled during the late steps of the viral cycle is largely unknown. Using super-resolution microscopy, we show that HIV-1 buds in areas of the plasma membrane with low cortical F-actin levels. We find that the cellular oxidoreductase MICAL1 locally depolymerizes actin at budding sites to promote HIV-1 budding and release. In the absence of MICAL1, F-actin abnormally remains at viral budding sites, incompletely budded viruses accumulate at the plasma membrane and viral release is impaired. Remarkably, normal viral release can be restored in MICAL1-depleted cells by inhibiting Arp2/3-dependent branched actin networks. Mechanistically, we find that MICAL1 directly disassembles branched-actin networks and controls the timely recruitment of the ESCRT scission machinery during viral budding. In addition, the MICAL1 activator Rab35 is recruited at budding sites, functions in the same pathway as MICAL1 and is also required for viral release. This work reveals a role for oxidoreduction in triggering local actin depolymerization to control HIV-1 budding, a mechanism that may be widely used by other viruses. The debranching activity of MICAL1 could be involved beyond viral budding in various other cellular functions requiring local plasma membrane deformation.

## INTRODUCTION

The plasma membrane undergoes continuous remodeling to ensure cellular events such as endocytosis, exocytosis, cell division, or viral budding. The egress of enveloped viruses from infected cells and the final step of cell division —cytokinetic abscission— require membrane deformation and topologically equivalent scission events mediated by the same cellular Endosomal Sorting Complexes Required for Transport (ESCRT) machinery (1-10). Whether other mechanisms are shared during cytokinetic abscission and viral budding remains largely unexplored.

HIV-1 budding involves coordinated interactions between viral proteins and the plasma membrane (11-13). The viral Gag polyprotein plays a crucial role by orchestrating virus assembly, budding and release. Gag polyproteins first bind to the plasma membrane, then cluster into PtdIns(4,5)P2-rich domains to generate viral particles in which viral RNA is packaged. Finally, Gag recruits the ESCRT machinery required for membrane scission and viral release from the infected cell.

Below the plasma membrane, the actin cortex forms a 200 nm-thick, dense and crosslinked meshwork of actin filaments (14-17). The actin cortex and its tight association with the plasma membrane imposes a significant challenge for membrane deformation and scission during the final steps of HIV-1 cycle. Several studies have reported an active role for actin dynamics in the assembly and budding of HIV-1 (18-22) while others argued against (23, 24). While the exact role of the actin cortex in the late steps of HIV-1 cycle remains debated (25), it is likely that membrane deformability and scission depend on actin dynamics at virus assembly and budding sites. In particular, the potential interplay between the actin cytoskeleton and the ESCRT-III machinery during budding is unknown.

During cytokinesis, the clearance of actin filaments (F-actin) from the intercellular bridge (ICB) connecting the two daughter cells is required for the scission of the plasma membrane and thus successful abscission (26-34). We previously demonstrated that the oxidoreductase MICAL1 —an enzyme that accelerates actin filament disassembly — is recruited and activated in the ICB by the small GTPase Rab35 to locally depolymerize actin filaments, thus promoting cytokinetic abscission (34). MICAL1 belongs to a conserved family of flavoenzyme oxidoreductases named MICAL (molecule interacting with CasL) that directly oxidize specific methionine residues in actin filaments, leading to their rapid disassembly (35-39). Mechanistically, actin disassembly by MICAL1 promotes ESCRT-III localization at the abscission site and thus membrane scission (34).

Here, we hypothesized that cortical actin filaments are similarly depolymerized by Rab35/MICAL1 during HIV-1 budding at the plasma membrane to facilitate the ESCRT-III-dependent release of viruses from infected cells. We found that HIV-1 budding preferentially occurs in regions of the plasma membrane with reduced F-actin levels. In contrast, F-actin remains at high levels at budding sites in absence of MICAL1, whose depletion induces an accumulation of incompletely budded viral particles at the plasma membrane and delays ESCRT-III recruitment at budding sites, leading to impaired HIV-1 release from infected cells. Finally, our results indicate that MICAL1 and its activator Rab35 function in the same pathway during HIV-1 budding and release. Altogether, this work reveals a role for oxidoreduction in retroviral budding, linking MICAL-1-mediated oxidation to local actin depolymerization at HIV-1 budding sites.

## RESULTS

### MICAL1 depletion reduces HIV-1 release from infected cells

We first investigated the role of MICAL1 in late stages of the HIV-1 cycle over one single viral cycle. To this aim, we silenced MICAL1 in HeLa cells using a specific small interfering RNA (siRNA) and then infected the cells with the HIV-1 strain NL4-3 pseudotyped with the vesicular stomatitis virus G glycoprotein (NL4-3-VSVG). Treatment with MICAL1 siRNAs led to an efficient depletion of MICAL1 in infected cells (*SI Appendix*, Fig. S1A). The day after infection, we exchanged the media to remove residual viral input and let cells produce viruses for 24 hours. Viral proteins both in infected cells and released in the supernatant were then analyzed by Western Blot (WB) using anti-Gag antibodies (Fig. 1A). The levels of Gag-p24 in the supernatant were reduced after MICAL1 depletion, with no decrease in the intracellular Gag precursor (Gag-p55) and Gag subproducts (Gag-p41 + Gag-p24) levels. We then quantified the levels of Gag-p24 in both the supernatant and infected cells by ELISA. MICAL1 depletion resulted in a 60 ± 16 % reduction of HIV-1 release (Fig. 1B). These results were confirmed with a second siRNA targeting MICAL1 (*SI Appendix*, Fig. S1B and S1C). We conclude that MICAL1 is required for efficient release of HIV-1 from infected cells.

**Figure 1:**
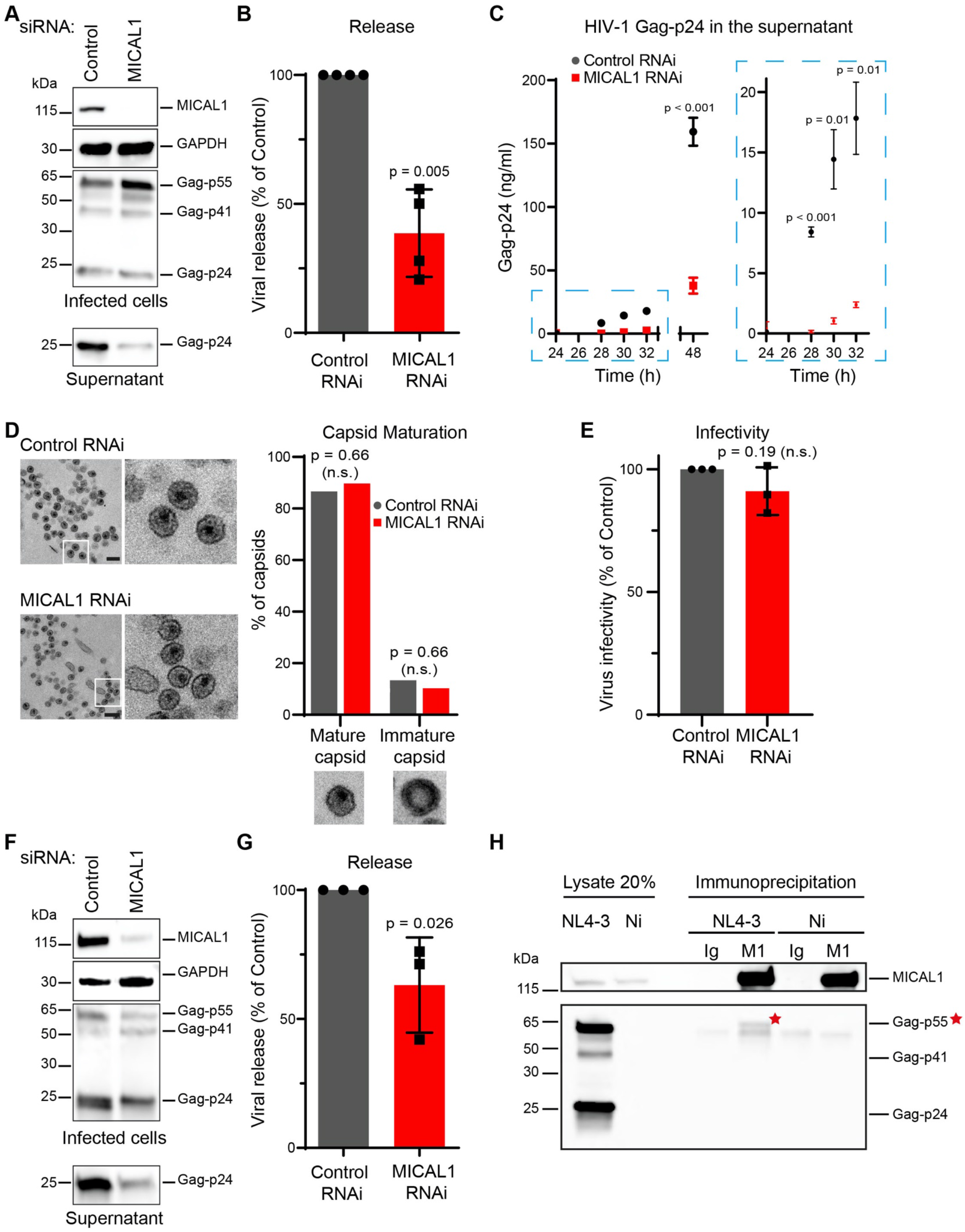
MICAL1 depletion reduces HIV-1 release from infected cells. **(A)** Western blot analysis of HIV-1 Gag products in infected HeLa cells treated with indicated siRNAs (panels Infected cells) and corresponding supernatants (panel Supernatant). Loading control: GAPDH. Equal volumes of cell and supernatant samples were loaded. This experiment was repeated at least 3 times independently with similar results. **(B)** Quantification by Gag-p24 ELISA of HIV-1 release in HeLa cells treated with either Control or MICAL1 siRNAs. Results were normalized to control RNAi conditions (set at 100 %). Error bars represent SD calculated from 4 independent experiments, each done in triplicate. Two-tailed unpaired Student’s t-test. **(C)** Quantification by ELISA of released Gag-p24 in supernatant of infected HeLa cells treated with either Control or MICAL1 siRNAs at indicated times after infection. Error bars represent SD calculated from one experiment done in triplicate. Two-tailed unpaired Student’s t-test. **(D)** Left panels: Transmission electron microscopy images of released viruses from infected HeLa cells treated with either Control or MICAL1 siRNAs. Scale bars, 200 nm. Right panel: Quantification of the proportion of mature vs. immature capsids. n = 232 (Control RNAi) and n = 165 (MICAL1 RNAi) released capsids. Fisher’s exact test. **(E)** The virus infectivity was scored by measuring beta-Galactosidase levels in infected HeLa P4C5 reporter cells treated with either Control or MICAL1 siRNAs. The beta-Galactosidase values were normalized to the amount of released Gag-p24. Results were normalized to control RNAi conditions (set at 100 %). Error bars represent SD calculated from 3 independent experiments. Two-tailed unpaired Student’s t-test. **(F)** Western blot analysis of HIV-1 Gag products in infected THP-1 cells treated with indicated siRNAs (panels Infected cells) and corresponding supernatants (panel Supernatant). Loading control: GAPDH. Equal volumes of cell and supernatant samples were loaded. This experiment was repeated at least 3 times independently with similar results. **(G)** Quantification by Gag-p24 ELISA of HIV-1 release in THP-1 cells treated with either Control or MICAL1 siRNAs. Results were normalized to control RNAi conditions (set at 100 %). Error bars represent SD calculated from 3 independent experiments, each done in triplicate. Two-tailed unpaired Student’s t-test. **(H)** Endogenous MICAL1 from infected (NL4-3) or non-infected (Ni) HeLa cells was and revealed with anti-MICAL1 antibodies (control immunoprecipitation = Rabbit IgG). Co-immunoprecipitated Gag-p55 (red star) was revealed with anti-Gag antibodies. The lower band corresponds to non-specific background also present in non-infected cells.

To determine when the release of the virus starts to be affected by the MICAL1 depletion, we monitored the kinetics of Gag-p24 release in the supernatant of infected cells. Twenty-four hours post-infection, we replaced the culture medium with fresh medium and collected viral supernatants at 4 – 6 – 8 and 24 hours and measured Gag-p24 by ELISA. At the end of the experiment, we collected infected cells and found that cell-associated Gag-p24 levels were unchanged in MICAL1-depleted cells, (*SI Appendix*, Fig. S1D). In contrast, the amounts of Gag-p24 in the supernatant were reduced in MICAL1-depleted cells at all time points (Fig. 1C), suggesting that HIV-1 release is impaired as soon as the infected cells start to produce viruses. Of note, MICAL1 depletion affected the release of both immature (Gag-p55) and mature (Gag-p24) as shown by WB analysis of purified viruses (*SI Appendix*, Fig. S1E).

We next investigated whether the maturation and the infectivity of the released viruses were affected by MICAL1 depletion. First, we analyzed virion maturation by transmission electron microscopy (TEM) and classified virions as either immature —characterized by discernible immature Gag shells— or mature —characterized by discernible conical or condensed cores (Fig. 1D). In control cells, 87 % of the released virions were mature (n= 232 virions). Similar results were obtained in MICAL1-depleted cells (90 % mature, n=165 virions) (Fig. 1D). We next used the HeLa P4C5 reporter cells to assess viral infectivity after infection with equal amounts of viral particles. We observed that infectivity of the released viruses was unchanged after MICAL1 depletion (Fig. 1E). Thus, MICAL1 depletion reduces HIV-1 release but does not perturb viral maturation nor subsequent infectivity.

To confirm the role of MICAL1 during viral release in a cell type naturally infected by HIV-1, we depleted MICAL1 in monocyte-derived macrophage THP-1 cells and infected them overnight with NL4-3-VSVG viruses. We then replaced the culture medium with fresh medium and collected the virus produced over the next 48 hours. We observed that the levels of Gag-p24 in the supernatant were reduced after MICAL1 depletion, as assessed by WB (Fig. 1F). We quantified the viral release by Gag-p24-ELISA and found that MICAL1 depletion in THP-1 cells resulted in a 37 ± 21 % reduction of HIV-1 release (Fig. 1G).

We finally investigated potential interactions between MICAL1 and Gag using immunoprecipitation assays. We infected HeLa cells with the NL4-3-VSVG virus, immunoprecipitated endogenous MICAL1 and found that it co-immunoprecipitated Gag-p55 (Fig. 1H). To confirm this interaction, we transfected HeLa cells with a NL4-3-Gag-GFP provirus, immunoprecipitated Gag-GFP and found that it co-immunoprecipitated endogenous MICAL1 (*SI Appendix*, Fig. S1F). These results indicate that MICAL1 and Gag form a complex and show that MICAL1 functions during viral budding and/or release.

### MICAL1 depletion induces accumulation of HIV-1 budding particles at the plasma membrane

Reduction of viral particle release can result from a defect during the budding process, leading to the accumulation of viruses at the plasma membrane of infected cells. We first analyzed the distribution of Gag and Env (viral envelope glycoprotein) structural proteins using spinning disk confocal microscopy 36 hours after NL4-3-VSVG infection in HeLa cells, both in the absence or presence of MICAL1. As expected, we observed a co-localization of the Gag and Env signals at the plasma membrane in both conditions (Fig. 2A and *SI Appendix*, Fig. S2A).

**Figure 2:**
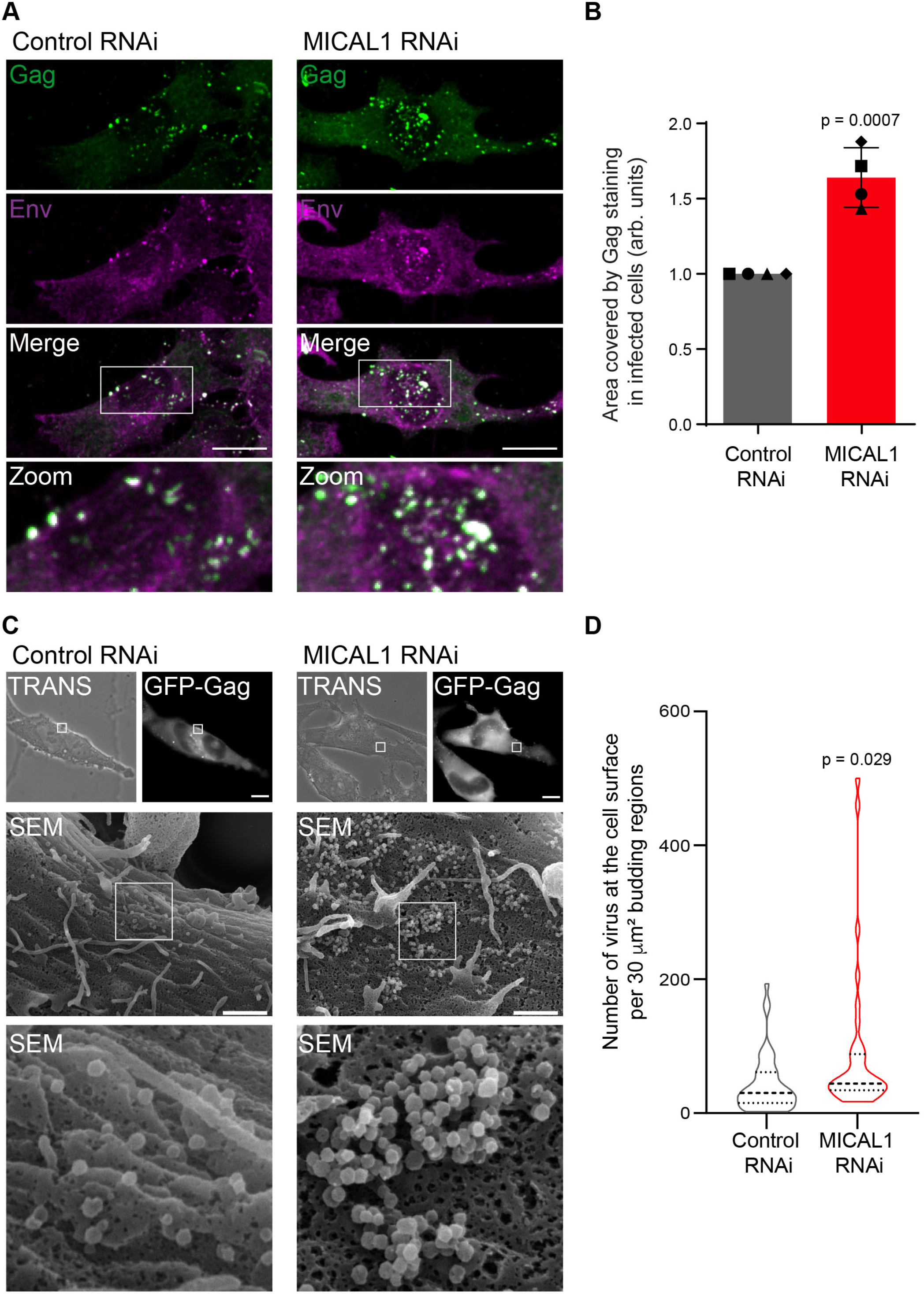
MICAL1 depletion induces accumulation of HIV-1 budding particles at the plasma membrane. **(A)** Z projection of spinning disk confocal images of infected HeLa cells treated with indicated siRNAs and labeled with anti-Gag (green) and anti-Env (magenta) antibodies. Scale bars, 10 μm. **(B)** Quantification of the area covered by Gag staining in infected cells (arbitrary units, arb. units). Results were normalized to control RNAi conditions (set at 1.0) Error bars represent SD calculated from 4 independent experiments. n = 145 (Control RNAi) and n = 154 (MICAL1 RNAi) cells analyzed. Two-tailed unpaired Student’s t-test. **(C)** Correlative light-scanning electron microscopy (SEM) of HeLa cells transfected with NL4-3-Gag-GFP provirus after control (left) or MICAL1 (right) depletion. Phase contrast (TRANS), fluorescent and SEM pictures with corresponding zooms of budding regions are presented. Scale bars, 10 μm for fluorescence pictures and 1 μm for SEM. **(D)** Quantification of the number of viruses at the cell surface per 30 μm^2^ budding regions (violin plot with median and interquartiles). n = 47 budding regions from 13 cells analyzed in Control RNAi conditions and n = 35 budding regions from 9 cells analyzed in MICAL1 RNAi conditions. Kolmogorov-Smirnov test.

Automatic detection of Gag-positive regions (*SI Appendix*, Fig. S2B top panels) revealed that MICAL1 depletion induced an increase in both the number and the surface of individual Gag-positive regions resulting in an increase of the total area covered by the Gag staining (Fig. 2B and *SI Appendix*, Fig. S2B bottom panels). These results indicate that MICAL1 depletion induces the formation of larger and more numerous Gag/Env regions at the plasma membrane, suggesting an accumulation of budding viruses at the cell surface.

This was confirmed by visualizing single viral particles on the surface of infected cells, using correlative light scanning electron microscopy (SEM) in HeLa cells infected with a NL4-3-Gag-GFP virus (Fig. 2C). We observed a 2-fold increase in the density of viruses at Gag-GFP-positive budding regions in MICAL1-depleted cells, as compared to control cells (Fig. 2D).

Altogether, these results show that MICAL1 depletion induces accumulation of budding viruses at the surface of infected cells, which likely accounts for the defective viral release observed in MICAL1-depleted cells.

### MICAL1 depletion impairs normal budding and delays ESCRT-III recruitment at budding sites

The accumulation of budding viruses in MICAL1-depleted cells suggests incomplete viral budding and/or defective scission. Incomplete viral budding results in the accumulation of budding events with a characteristic “half-moon” shape (40, 41). Conversely, a defect in viral particle scission leads to the formation of a typical “lollipop” shape budding particles (42). Analysis of TEM images from MICAL1-depleted cells infected with NL4-3-VSVG revealed an accumulation of incompletely budded viruses at the plasma membrane (Fig. 3A). Budding virions in MICAL1-depleted cells displayed decreased height (72 ± 33 nm, n= 134), as compared to control cells (99 ± 25 nm, n= 132), while the widths were similar (133 ± 25 nm and 129 ± 22 nm in control and in MICAL1-depleted cells, respectively) (Fig. 3B). Consequently, the height-to-width ratio decreased by 32 % in MICAL1-depleted cells (Fig. 3B). Thus, MICAL1 depletion leads to the accumulation of incomplete budded particles at the plasma membrane.

**Figure 3:**
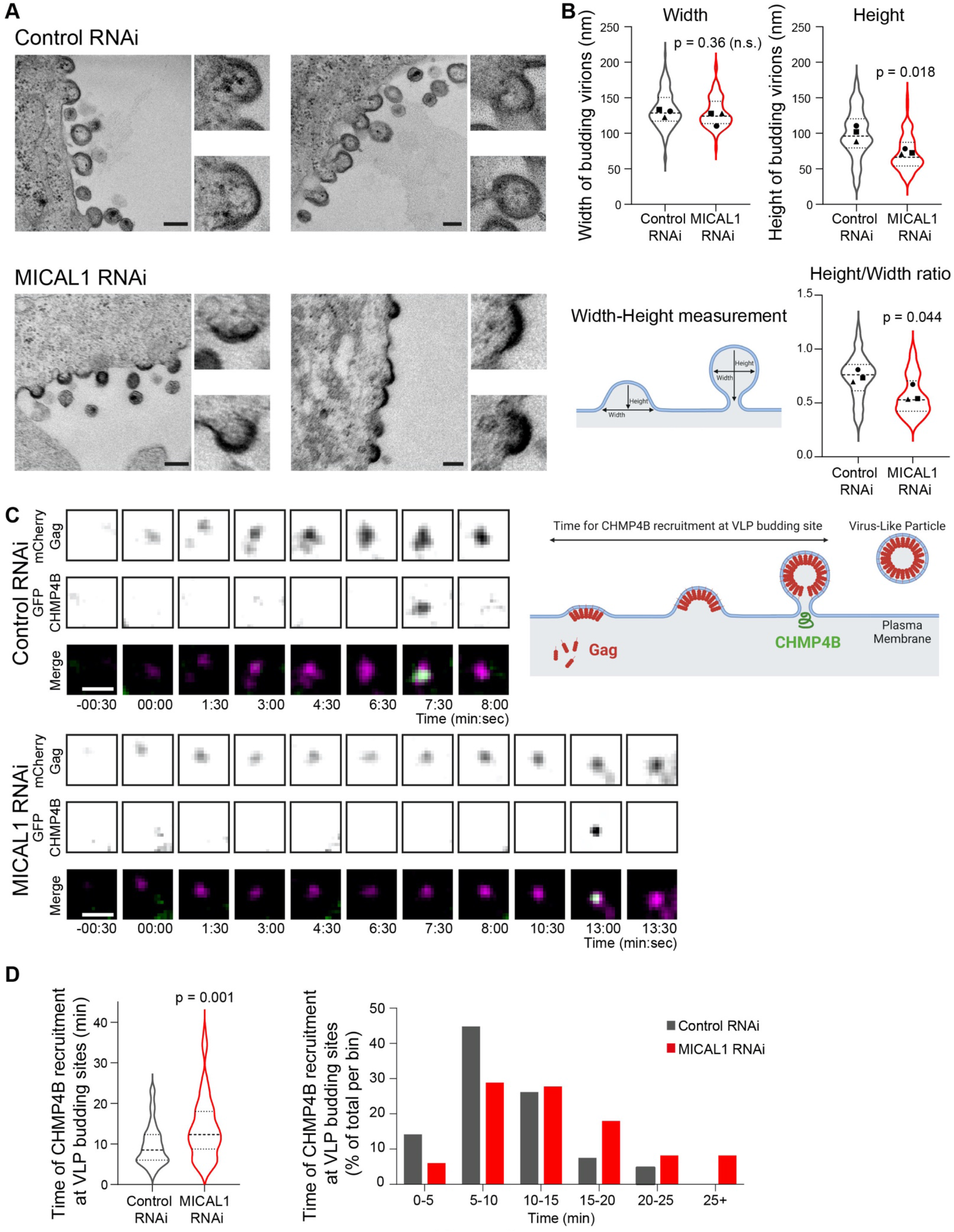
MICAL1 depletion impairs normal budding and delays ESCRT-III recruitment at budding sites. **(A)** Transmission electron microscopy (TEM) of infected HeLa cells treated with either Control of MICAL1 siRNAs. Scale bars, 200 nm. **(B)** Width, Height and Height/Width ratio of budding virions in infected HeLa cells treated with either Control or MICAL1 siRNAs. Mean of 3 independent experiments (violin plot with median and interquartiles). n = 132 (Control RNAi) and n = 134 (MICAL1 RNAi) buds. Two-tailed unpaired Student’s t-test. **(C)** Time-lapse images (one image every 30 seconds for 45 minutes) using TIRF microscopy of HeLa cells stably expressing GFP-CHMP4B (green) and transfected with plasmid encoding Gag-mCherry (magenta) after treatment with either Control or MICAL1 siRNAs. Scale bars, 1 μm. **(D)** Quantification of the time of CHMP4B recruitment at VLP budding sites. Left: violin plot with median and interquartiles. Right: histograms of distribution of CHMP4B recruitment times. n = 75 VLPs from 8 cells and 92 VLPs from 8 cells in Control RNAi and MICAL1 RNAi conditions, respectively. N = 3 independent experiments. Kolmogorov-Smirnov test.

We next investigated whether MICAL1 depletion impaired the recruitment of the ESCRT-III protein CHMP4B at budding sites. First, infected control and MICAL1-depleted cells were stained with anti-Gag and anti-CHMP4B antibodies and analyzed by spinning disk microscopy to quantify the percentage of colocalization between these two proteins (*SI Appendix*, Fig. S2C). In Gag-positive regions of control cells, 40% of the Gag-positive pixels colocalized with CHMP4B. In the absence of MICAL1, this percentage was significantly reduced to 26% (*SI Appendix*, Fig. S2C). This suggests that MICAL1 promotes CHMP4B recruitment to the budding regions. We then performed total internal reflection fluorescence (TIRF) microscopy in HeLa cells stably expressing GFP-tagged CHMP4B (43) and transfected with a plasmid encoding a carboxy-terminally tagged Gag (Gag-mCherry), leading to the formation of virus-like particles (VLPs) that bud at the plasma membrane (44). By tracking Gag and CHMP4B kinetics with time lapse microscopy, we determined the time for CHMP4B recruitment at VLP budding sites (Fig. 3C and *SI Appendix*, Fig. S2D). In the representative examples presented in Figure 3C, the mCherry signal gradually accumulates until GFP-CHMP4B is recruited at the budding site after 7 min 30 sec in control cell vs. 13 min in MICAL1-depleted cell (T0 being the first time point where we could detect a mCherry-Gag spot at the plasma membrane). The analysis of 75-92 budding events where CHMP4B was recruited showed that CHMP4B recruitment occurred 9.6 ± 5 minutes after Gag arrival in control cells (Fig. 3D), similarly to what has been previously observed (45). In contrast, the time necessary to recruit CHMP4B following Gag appearance was increased by 45 % to 13.9 ± 7 minutes in MICAL1-depleted cells (Fig. 3D). Thus, CHMP4B recruitment at the budding site is delayed in MICAL1-depleted cells.

### MICAL1 depletion impairs F-actin clearance at HIV-1 budding sites

Since MICAL1 binds and disassembles actin filaments, we hypothesized that MICAL1 depletion could affect F-actin levels at HIV-1 budding sites. Using spinning disk confocal microscopy, we observed that, in control cells, F-actin levels —visualized by fluorescent phalloidin— were reduced in Gag-positive regions of the plasma membrane, compared to adjacent Gag-negative regions (Fig. 4A left panels and 4B). In contrast, in MICAL1-depleted cells, F-actin levels in Gag-positive regions remained has high as in Gag-negative regions (Fig. 4A right panels and 4B). We next classified the budding regions into three categories based on F-actin intensity in Gag-positive regions compared to adjacent Gag-negative regions: (1) reduced F-actin in Gag-positive regions (at least 30% reduced intensity), (2) increased F-actin in Gag-positive regions (at least 30% increased intensity), and (3) no significant differences in F-actin levels (between 30% reduced intensity and 30% increased intensity). In control cells, 60 % of budding regions localized in areas of reduced F-actin while in MICAL1-deficient cells, this was observed in only 39 % of budding regions (Fig. 4C). Furthermore, in MICAL1-depleted cells, there was a significant increase in budding regions localized in areas where F-actin levels were similar to nearby regions (Fig. 4C). We confirmed in Jurkat T cells and activated primary CD4 T cells infected with the NL4-3-VSVG virus that HIV-1 budding occurred mainly in the regions of the plasma membrane where cortical actin was reduced (in 60.8% in Jurkat T cells and in 57.4% in primary CD4 T cells) (*SI Appendix*, Fig. S3A and S3B). This is consistent with a recent report showing that Gag assembly occurs in less dense in F-actin area at the T cell plasma membrane (22).

**Figure 4:**
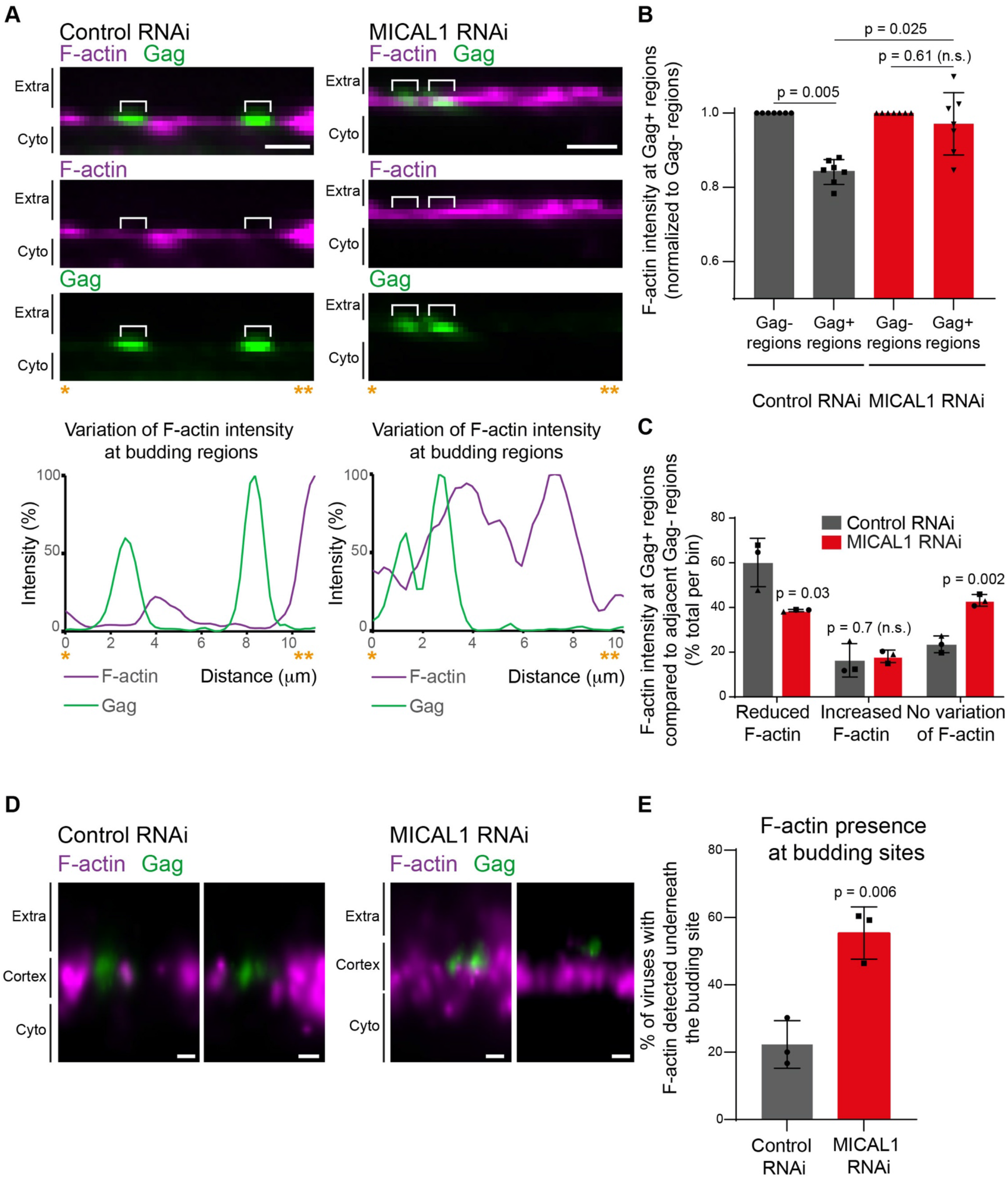
MICAL1 depletion impairs F-actin clearance at HIV-1 budding sites. **(A)** Top panels: Spinning disk confocal images of infected HeLa cells treated with either Control (left) or MICAL1 (right) siRNAs and labeled with anti-Gag (green) antibody and fluorescent phalloidin (magenta). Extra: Extracellular space. Cyto: cytoplasm. Bottom panels: Plot profile of the Gag and Phalloidin intensity of corresponding images. Scale bars, 2 μm. **(B)** Quantification of F-actin intensity (using fluorescent phalloidin) at Gag-positive (Gag+) and Gag-negative (Gag-) regions (normalized to Gag-regions), in Control and MICAL1-depleted HeLa cells (n = 132 and n = 154 Gag+ budding regions from 7 cells analyzed in Control RNAi and MICAL1 RNAi conditions, respectively). Error bars represent SD. 2 way-ANOVA multiple comparisons. **(C)** Quantification of the variations of F-actin intensity in Gag+ regions compared to adjacent Gag-regions. n = 176 Gag+ regions and n = 196 Gag+ regions from 12 cells analyzed in Control and MICAL1 RNAi conditions, respectively. Error bars represent SD calculated from 3 independent experiments. Two-tailed unpaired Student’s t-test. **(D)** 3D-STORM images of infected HeLa cells treated with either Control (left) or MICAL1 (right) siRNAs and labeled with anti-Gag antibody and fluorescent phalloidin. Scale bars, 200 nm. **(E)** Quantification of F-actin presence at budding sites using 3D-STORM images. n = 74 and n = 98 budding viruses from 3 cells analyzed in Control RNAi and MICAL1 RNAi conditions, respectively. Error bars represent SD. Two-tailed unpaired Student’s t-test.

To visualize the cortex underneath budding sites at high resolution, we turned to super-resolution microscopy using two-color three-dimension stochastic optical reconstruction microscopy (3D-STORM). We did not detect F-actin beneath most budding viruses in control cells (Fig. 4D left panels and 4E). In contrast, F-actin was present beneath budding viruses after MICAL1 depletion (Fig. 4D right panels and 4E).

Altogether, these results indicate that HIV-1 buds preferentially in areas of the plasma membrane with relatively low cortical F-actin levels, and reveal that MICAL1 is key for locally clearing cortical actin at budding regions.

### Inhibiting branched-actin nucleation restores normal HIV-1 release in MICAL1-depleted cells

The results above imply that decreasing F-actin levels at the cell cortex is required for efficient viral budding and release. We thus investigated whether the defects in viral release found in MICAL1-depleted cells could be rescued by inhibiting F-actin nucleation. To do so, we treated infected cells with the Arp2/3 inhibitor CK666 (75 μM) to reduce the formation of branched F-actin networks. We first checked that the CK666 treatment at this concentration did not alter MICAL1 depletion (*SI Appendix*, Fig. S4A), did not impact cell viability (*SI Appendix*, Fig. S4B) and did not significantly affect HIV-1 release in control cells (Fig. 5A). Remarkably, CK666 treatment in MICAL1-depleted cells restored HIV-1 release (Fig. 5A) and the total area of budding regions (*SI Appendix*, Fig. S4C and quantified in Fig. 5B) to levels comparable to those observed in control, non-depleted cells. Thus, reducing branched-actin levels restores normal HIV-1 release in MICAL1-depleted cells. Furthermore, these results indicate that the abnormal persistence of F-actin at HIV-1 budding sites resulting from MICAL1 depletion is likely responsible for the defects in virus release.

**Figure 5:**
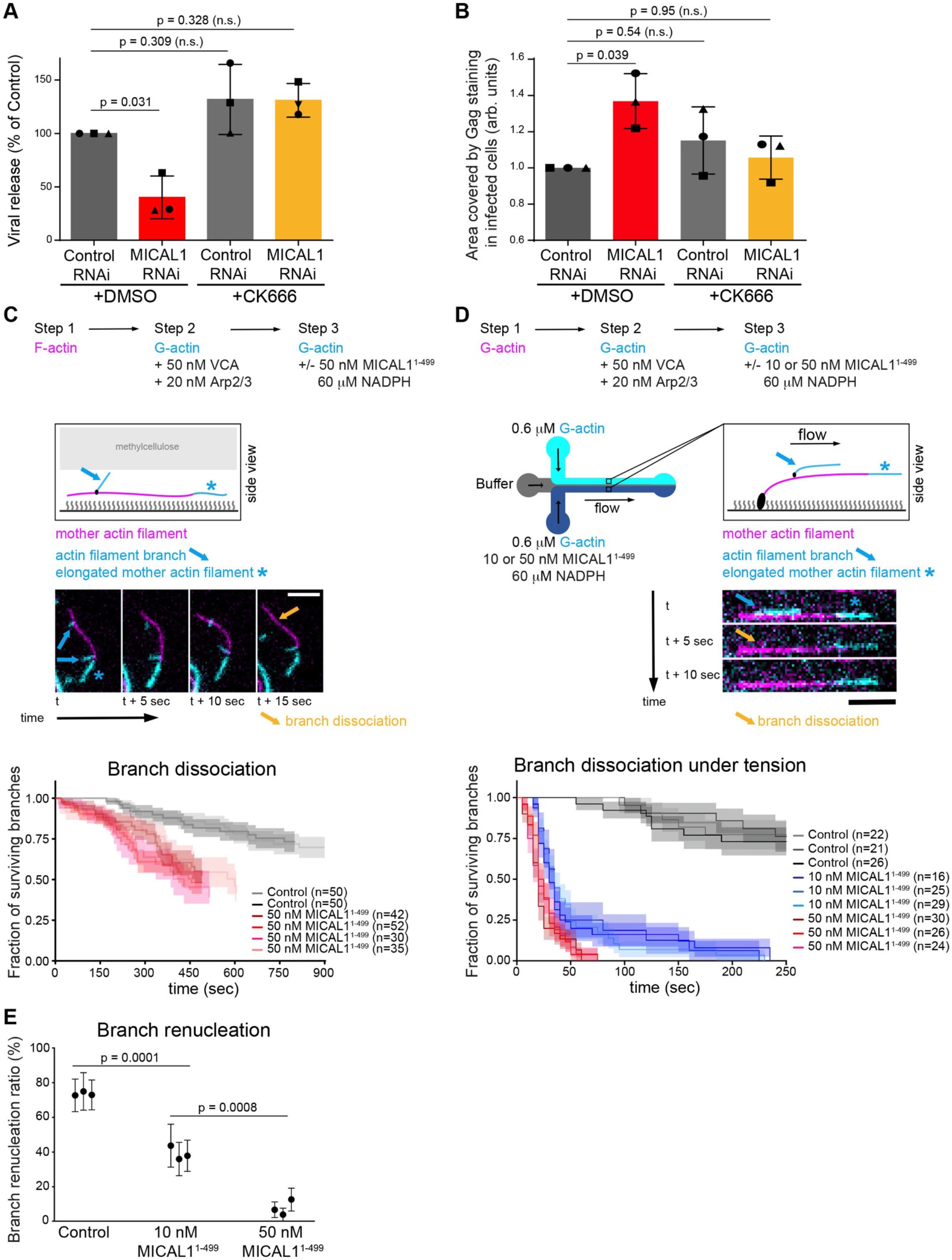
Inhibiting branched-actin nucleation restores normal HIV-1 release in MICAL1-depleted cells and MICAL1 exhibits debranching activity. **(A)** Quantification by Gag-p24 ELISA of HIV-1 release in Control and MICAL1-depleted HeLa cells, treated with either DMSO or CK666 (75 μM). Results were normalized to control RNAi treated with DMSO conditions (set at 100 %). Error bars represent SD calculated from 3 independent experiments, each done in triplicate. One-way ANOVA multiple comparisons. **(B)** Quantification of the area covered by Gag staining in infected cells (arbitrary units, arb.units.). Error bars represent SD calculated from 3 independent experiments. n = 185, 188, 206 and 185 cells analyzed for Control RNAi + DMSO, Control RNAi + CK666, MICAL1 RNAi + DMSO and MICAL1 RNAi + CK666 conditions, respectively. One-way ANOVA multiple comparisons. **(C)** Top panel: schematic of branching and debranching experiment in an in vitro reconstitution assay using an open microchamber. Middle panel: fluorescence microscope image sequence, showing the dissociation (yellow arrow) of an actin filament branch (cyan + blue arrow) from its mother filament (magenta, and cyan + blue star). The time interval between images is 5 seconds. Filaments were monitored in the presence of 0.3 μM 10% ATTO-488 labeled G-actin (cyan) —causing all barbed ends to elongate—, with or without 50 nM MICAL1^1-499^ (active catalytic domain) and 60 μM NADPH. Scale bar, 5 μm. Bottom panel: fraction of remaining (undissociated) actin filament branches as a function of time, in the presence or absence of MICAL1^1-499^ and NADPH. The shaded areas show the 65% confidence intervals. Between control (pooled repeats) and 50 nM (pooled repeats), p<0.0001 (log-rank test). **(D)** Top panels: schematic of the debranching experiment in an in vitro reconstitution assay using a microfluidics chamber. Filaments are exposed to different conditions (with or without MICAL1^1-499^ and NADPH) side by side, in the same chamber. Middle panel: the microscope image sequence shows the dissociation (yellow arrow) of an actin filament branch (cyan + blue arrow) from its mother filament (magenta, and cyan + blue star). The time interval between images is 5 seconds. Scale bar, 5 μm. Bottom panel: Fraction of remaining (undissociated) actin filament branches as a function of time, under indicated conditions. Each curve shows the surviving fraction of different populations of n branches over time. In all these experiments, the average force on branch junctions was 0.32 (+/-0.03, SD) pN at t=0 and increased at a rate of 0.00056 (+/-0.00011, SD) pN/s as the branches elongated over time. The shaded areas show the 65% confidence intervals. Between different concentrations (pooled repeats), p<0.0001 (log-rank test). **(E)** Percentage of branches that renucleate after dissociating, in the presence of 60 µM NADPH and different concentrations of MICAL1^1-499^. Each data point corresponds to one experiment, monitoring, from left to right, n= 22,16, 26, 16, 25, 29, 30, 26, 24 dissociated branches (same filament populations as in D). Error bars show binomial standard deviations.

### MICAL1 exhibits debranching activity

The CK666 rescue experiments also suggest that MICAL1 promotes the disassembly of branched actin networks. Yet, such a debranching activity of MICAL1, beyond its known function in actin disassembly, has not been reported so far. We thus tested in vitro whether MICAL1 can promote the dissociation of F-actin branches, using fluorescence microscopy and purified proteins to monitor individual events in controlled conditions. To specifically characterize this new activity on actin branches, we prevented barbed end depolymerization by ensuring sufficient levels of non-oxidized G-actin. First, we monitored the dissociation of actin filament branches in the absence of mechanical forces. We found that debranching occurred more rapidly in the presence of the monooxygenase catalytic domain of MICAL1 (MICAL1^1-499^) and its coenzyme nicotinamide adenine dinucleotide phosphate (NADPH) (Fig. 5C). Since oxidation of F-actin by MICAL1 can favor the severing of filaments by weakening intrafilamentary bonds (46), we verified that the observed debranching was not due to severing of the oxidized filaments in our experiments (*SI Appendix*, Fig. S4D). As cortical F-actin branches are likely exposed to mechanical stress at the cell cortex, we then asked whether MICAL1 would also promote debranching under pulling forces. We thus monitored branch dissociation using microfluidics (47), which enables the application of a controlled pulling force to branched filaments (48). As previously reported (48, 49), the application of mechanical force alone accelerated branch dissociation (Fig. 5D, grey curves). In addition, the presence of MICAL1 strongly accelerated debranching, in a dose-dependent manner (Fig. 5D, red and blue curves). The presence of purified Gag did not modify the debranching activity of MICAL1 in vitro (*SI Appendix*, Fig. S4E), consistent with the absence of direct interaction between Gag and MICAL1 in 2-hybrid assays (*SI Appendix*, Fig. S4F). Since we recently discovered that the Arp2/3 complex remains bound to the mother filament and nucleates a new branch following most branch dissociation events (48), we also monitored the impact of MICAL1 on branch renucleation in microfluidics experiments (Fig. 5E). We found that the presence of MICAL1 efficiently prevented the renucleation of branches. Thus, our results reveal that MICAL1 has a debranching activity and regulates the dynamics of branched actin networks.

### MICAL1 and its activator the GTPase Rab35 function in the same pathway in HIV-1 release

MICAL1 enzymatic activity is tightly regulated in cells. In cytokinesis, MICAL1 is recruited and activated at the intercellular bridge by the small GTPase Rab35 to locally depolymerize actin filaments (34). We thus investigated the potential role of Rab35 in HIV-1 release and, if so, whether it acts in the same pathway as MICAL1. We silenced Rab35 alone or together with MICAL1 using specific siRNAs in infected HeLa cells and measured viral release by WB (Fig. 6A). Depletion of Rab35, alone or in combination with MICAL1, reduced released Gag-p24, as shown by WB quantifications (Fig. 6A and *SI Appendix*, Fig. S5A). Quantifications by Gag-p24-ELISA showed similar release defects after either MICAL1 depletion or Rab35 depletion (Fig. 6B). Importantly, no additive effect on viral release was observed when MICAL1 and Rab35 were co-depleted (Fig. 6B and *SI Appendix*, Fig. S5A). Furthermore, as in MICAL1-depleted cells, the total area covered by budding regions was increased in Rab35-depleted cells (*SI Appendix*, Fig. S5B and quantified in Fig. 6C). Compared to either Rab35 or MICAL1 depletion alone, no further increase in the total area covered by budding regions was observed when Rab35 and MICAL1 were depleted together (*SI Appendix*, Fig. S5B and Fig. 6C). These results are consistent with the idea that MICAL1 and its activator Rab35 functions in the same pathway in HIV-1 release.

**Figure 6:**
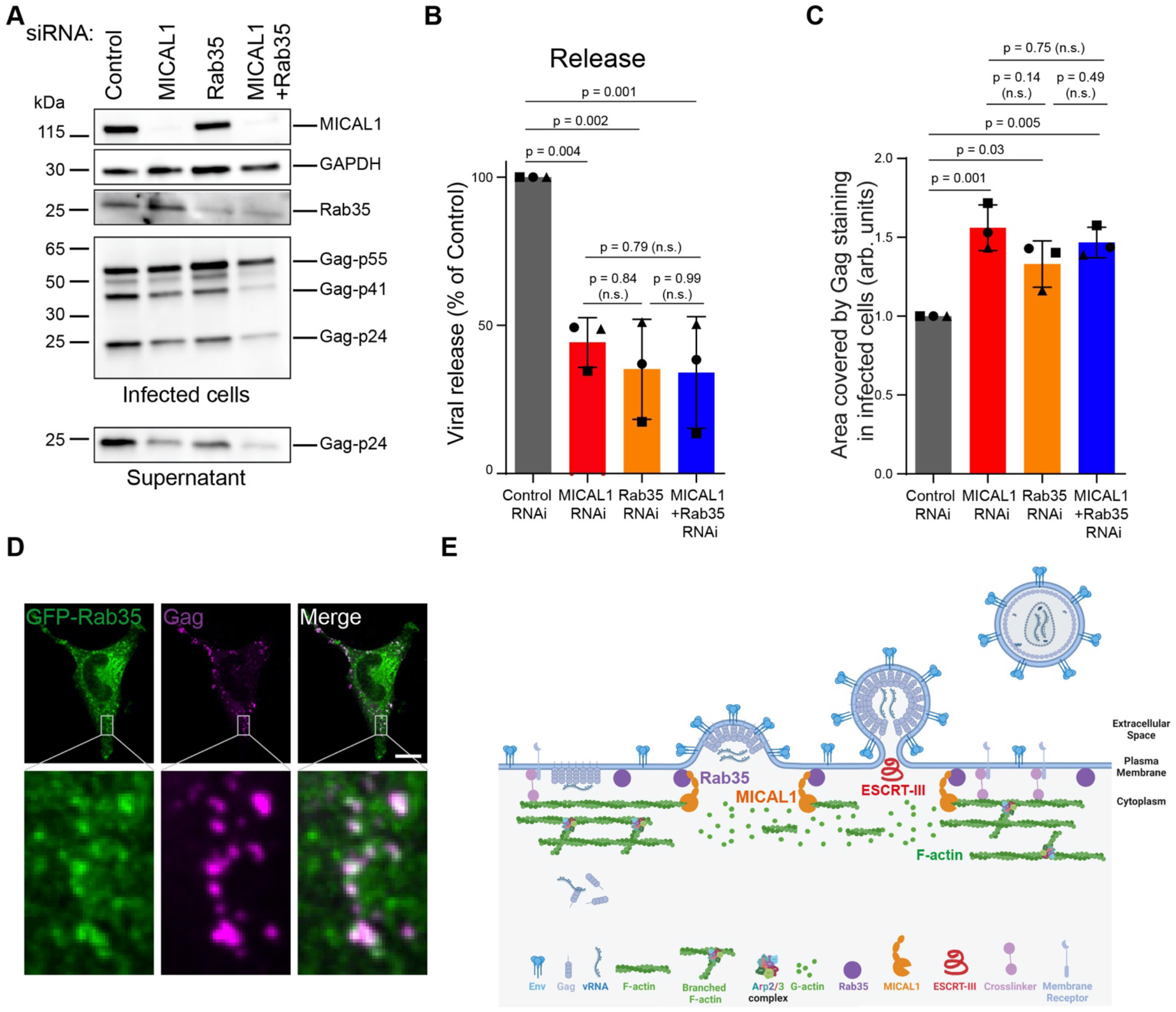
MICAL1 and its activator the GTPase Rab35 act in the same pathway in HIV-1 release. **(A)** Western blot analysis of HIV-1 Gag products in infected HeLa cells treated with indicated siRNAs (panels Infected cells) and corresponding supernatants (panel Supernatant). Loading control: GAPDH. Equal volumes of cell and supernatant samples were loaded. This experiment was repeated at least 3 times independently with similar results. **(B)** Quantification by Gag-p24 ELISA of HIV-1 release in infected HeLa cells treated with either Control, MICAL1, Rab35 or MICAL1 + Rab35 siRNAs. Results were normalized to control RNAi conditions (set at 100 %). Error bars represent SD calculated from 3 independent experiments, each done in triplicate. One-way ANOVA multiple comparisons. **(C)** Quantification of the area covered by Gag staining in infected cells (arbitrary units, arb.units.). Error bars represent SD calculated from 3 independent experiments. n = 126, 133, 122 and 131 cells analyzed for Control RNAi, MICAL1 RNAi, Rab35 RNAi and MICAL1 + Rab35 RNAi conditions, respectively. One-way ANOVA multiple comparisons. **(D)** Spinning disk confocal images of genome edited HeLa cells that express endogenous Rab35 tagged with GFP and stained with anti-Gag (magenta) antibody and GFP-Booster nanobody (green) after infection. Scale bars, 10 μm. **(E)** Model for F-actin clearance by MICAL1 at budding site during HIV-1 budding. We propose that Rab35 at HIV-1 budding sites activates the redox enzyme MICAL1, which locally disassembles cortical F-actin through its depolymerizing and debranching activities. This is required for complete budding and normal ESCRT-III recruitment, and thereby promotes viral release.

We next investigated whether Rab35 localized at HIV-1 budding regions. Using spinning disk confocal microscopy in a cell line where Rab35 is tagged with GFP at the endogenous locus (50), we detected the presence of Rab35 in 90 % of the budding regions (Fig. 6D, n = 277 budding regions in 9 infected cells, and *SI Appendix*, Fig. S5C). As a control, we used the GTPase Rab11 that does not interact with MICAL1 (51) and did not detect it at budding regions (*SI Appendix*, Fig. S5D). Altogether, we conclude that Rab35 localizes at viral budding sites, and we propose that Rab35 functions at this location to allow proper HIV-1 budding and release.

## DISCUSSION

Here, we report that the enzyme MICAL1 controls F-actin clearance at budding sites to promote HIV-1 budding and release (Fig. 6E). In the absence of MICAL1, F-actin remains underneath budding sites, CHMP4B recruitment is delayed, virus particles accumulate at the plasma membrane and HIV-1 release is reduced. Mechanistically, MICAL1 depletion affects the budding process, since incompletely budded particles accumulate at the plasma membrane in the absence of MICAL1.

HIV-1 assembles and buds from areas of the plasma membrane with reduced F-actin levels ((22) and this study). How could the abnormal presence of F-actin at budding sites observed upon MICAL1 depletion impact viral budding? One possibility is that F-actin clearance by MICAL1 promotes full Gag-mediated membrane deformation up to the point enabling the recruitment of ESCRT-III. In non-infected cells, membrane-cytoskeleton interactions affect plasma membrane mechanics and tension (15, 17, 52). In particular, proteins linking the plasma membrane to the actin cortex, such as Ezrin, Radixin, Moesin (ERMs), were shown to control membrane deformation (14) and have been implicated in HIV-1 cycle (53-55). High levels of F-actin at budding sites in the absence of MICAL1 may thus enhance ERM localization at these sites thereby preventing membrane deformation. Alternatively, these abnormal actin levels might impact on the localization of factors that play an active role in membrane deformation, such as the BAR-domain protein IRSp53 and Angiomotin (AMOT) (40, 41). Interestingly, the depletion of either IRSp53 or AMOT induces viral bud arrest at half completion, in a “half-moon” shape (40, 41), as observed upon MICAL1 depletion. It is thus possible that MICAL1 acts together with IRSp53/AMOT or in parallel in the same step of viral budding.

We found that MICAL1 depletion reduces HIV-1 release in two adherent cell lines, HeLa cells and monocyte-derived macrophages THP-1. However, we did not observe release defects in T lymphocytes upon MICAL1 depletion. Nevertheless, we found that budding viruses are present mainly in plasma membrane areas where F-actin level is reduced in both Jurkat T cells and primary activated CD4+ T cells, similarly to HeLa cells. One possibility is that MICAL1 requirement for actin disassembly at budding sites is different in adherent vs. non-adherent cells. Alternatively, the MICAL family comprises three members, MICAL1, MICAL2 and MICAL3, and have all been shown to induce F-actin depolymerization in vitro (36, 56). These enzymes could thus play a redundant role in viral budding in infected T-cells.

The fact that we were able to rescue the release defects in MICAL1-depleted cells by inhibiting Arp2/3 suggest that MICAL1 promotes the disassembly of branched actin during viral budding. Thus far, it is well established that MICAL1-dependent oxidation of linear actin filaments can induce their rapid depolymerization in vitro (34, 57-59). Here, we discovered that MICAL1, in addition, has a debranching activity by inducing the dissociation of actin filament branches from their mother filament. Our in vitro assays also revealed MICAL1’s ability to prevent the renucleation of dissociated branches, which is likely to play a role in situations of rapid filament turnover, as in the cell cortex. Recently, MICAL2 was found to promote the disassembly of the branched actin tails that propel vaccinia viruses within cells (60). It was shown that MICAL2 can oxidize Arp3B (60), but whether MICAL2 has a direct debranching activity has not been investigated. Since the catalytic domains of MICAL proteins are similar, it is likely that the debranching activity of MICAL1 that we observed in vitro is the consequence of the direct oxidation of the Arp2/3 complex. Further investigations are required to determine whether the oxidation of actin subunits in contact with the Arp2/3 complex at the branch junction also plays a role in actin branch dissociation.

A recent study reported that branched actin disassembly plays an early role during viral assembly (22). Indeed, infected T cells treated with CK666 favored the initial formation of Gag clusters at the plasma membrane, thereby increasing particle production. Consistently, in vitro experiments indicate that low F-actin density promote the initiation of Gag clustering on model membranes (22). It was further shown that Gag recruits the Arp2/3 inhibitory protein Arpin during viral assembly (22). An interesting possibility is that Arpin and MICAL1 function successively in the different steps of viral particle formation.

Our results indicate that MICAL1 acts locally at budding sites to depolymerize F-actin. However, we were unable to detect MICAL1 at budding sites using confocal spinning disk microscopy. This is likely due to the low amounts of MICAL1 present at the budding sites, that would require more sensitive approaches to be detected. Super-resolution microscopy, such as single-molecule localization microscopy (SMLN), could help for localizing MICAL1 at budding sites, as it has been successfully used for IRSp53 (41). Another possibility is that MICAL1 recruitment is too dynamic to be captured in fixed samples. Although technically challenging, live-cell microscopy on cells infected with tagged versions of the virus could be used to observe transient recruitment of MICAL1 at budding sites.

Besides being recruited at the right place at the right time, MICAL1 must be activated. Since it exhibits a strong actin depolymerizing activity, MICAL1 exists in an auto-inhibited conformation and either Plexin receptors or Rab GTPase binding to its C-terminal region releases this inhibition (57, 61-65). We previously demonstrated that the GTPase Rab35 interacts with this region, activates the enzymatic activity of MICAL1 in vitro and determines its localization at the ICB during cytokinesis (34). During viral assembly, Gag creates its own lipid nanodomains enriched in PtdIns(4,5)P2 at the plasma membrane (66-70). Interestingly, Rab35 contains an evolutionarily conserved polybasic C-terminal tail which facilitates its interaction with PtdIns(4,5)P2 and PtdIns(3,4,5)P3 at the plasma membrane (71, 72). In addition, Rab35 is incorporated into virions (73). However, its potential role in assembly, budding or release has not been investigated. Here, we show that Rab35 localizes at budding sites and promotes HIV-1 release. Our data further suggest that Rab35 and MICAL1 act in the same pathway in viral release. Thus, we propose that the Rab35/MICAL1 pathway, which locally clears F-actin at the plasma membrane, is used both for cytokinesis and viral budding.

In conclusion, we report a role of actin oxidation by MICAL1 in the late steps of the HIV-1 cycle and reveal the first connection between oxidoreduction and viral budding. This raises the possibility that other enveloped viruses use similar MICAL-dependent strategies to remodel the cortical actin in order to promote viral budding and release.

## MATERIALS AND METHODS

### Cell cultures

HeLa CCL-2 (ATCC) cell line, HeLa P4C5 cells (NIH AIDS Research and Reference Reagent Program), HeLa CHMP4B-GFP cell line (43) and GFP-Rab35 TALEN-edited HeLa cell line (50) were grown in Dulbecco’s Modified Eagle Medium (DMEM) GlutaMax (31966; Gibco, Invitrogen Life Technologies) supplemented with 10 % fetal bovine serum and penicillin–streptomycin (Gibco) at 50 ng/mL in 5 % CO_2_ at 37 °C. Jurkat T (ATCC) cells were grown in RPMI 1640 GlutaMax medium (61870010; Gibco, Invitrogen Life Technologies) supplemented with 10 % fetal bovine serum, 1× penicillin–streptomycin (Gibco) 50 ng/mL. Primary T CD4 cells were isolated from PBMCs obtained from healthy donors (Etablissement Français du Sang, EFS, Rungis) through positive selection using CD4+ T cell microbeads (Miltenyi 130-097-048). For T cell stimulation with PHA, the purified CD4+ T cells, maintained at a concentration of 5 × 10^6^ cells/mL in Roswell Park Memorial Institute (RPMI) medium, were treated overnight with PHA (1 μg/mL; Oxoid, Thermo Fisher). The following day, the cells were diluted 1:10 in interleukin 2 (IL-2)-containing RPMI (100 IU/mL; R&D Systems) supplemented with 10 % FBS and penicillin–streptomycin (RPMI-IL-2) and kept in culture until use. THP-1 (ATCC TIB-202) cells were grown in RPMI 1640 GlutaMax medium (61870010; Gibco, Invitrogen Life Technologies) supplemented with 10 % fetal bovine serum, penicillin–streptomycin (Gibco) at 50 ng/mL.

### Antibodies and plasmids

The following antibodies were used for western blot experiments: rabbit anti-MICAL1 (1:500, Proteintech Europe 14818-1-AP), rabbit anti-Rab35 (1:500, Proteintech Europe 11329-2-AP), mouse anti-GAPDH (1:8000 Proteintech Europe 60004-1-Ig), rabbit anti-Gag (NIH ARP-4250), mouse anti-Gag (NIH ARP-3537 183-H12-5C), mouse anti-ALIX (1:1000, Santa Cruz Biotechnology sc-271975) and mouse anti-GFP (1:2000 Proteintech Europe 66002-1-Ig). The following antibodies were used for immunofluorescence experiments: rabbit anti-Gag (NIH ARP-4250), mouse anti-Env (clone 110H – Institut Pasteur), rabbit anti-CHMP4B (13683-1-AP – Proteintech). The following secondary antibodies were used: Dylight Alexa 488- and Cy3- and Cy5-conjugated secondary antibodies (Jackson Laboratories) were diluted 1:500. Phalloidin staining (1:2000, Invitrogen). DAPI staining (0.5 mg/mL, Serva). GFP-Booster Alexa Fluor™ 488 (Proteintech ChromoTek). The plasmid encoding HIV-1 Gag-mCherry was a gift from Gregory Melikyan (Addgene plasmid # 85390 ; http://n2t.net/addgene:85390 ; RRID:Addgene_85390) (74). The plasmid encoding mCherry-Rab11 was a kind gift from Alexandre Gidon.

### siRNA transfections

For silencing experiments, HeLa cells were transfected with 25 nM siRNAs (siControl, siMICAL1, siMICAL1#2, siRab35) for 5 days using Lipofectamine RNAiMAX (Invitrogen), following the manufacturer’s instructions.

For silencing experiments, THP-1 cells were transfected with 50 nM siRNAs using Lipofectamine RNAiMAX. The day after, phorbol 12-myristate 13-acetate (PMA, Sigma) at 167 ng/mL were added to the culture medium to differentiate THP-1 cells in macrophages. Forty-eight hours later, a second siRNA transfection was done using Hiperfect (Qiagen). siRNAs against Luciferase (used as control, 5′CGUACGCGGAAUACUUCGA[dU][dU]3′), MICAL1 (5′GAGUCCACGUCUCCGAUUU[dU][dU]3′) (34), MICAL1#2 (5′CUCGGUGCUAAGAAGUUCU[dU][dU]3′) (75) and Rab35 (5′GCUCACGAAGAACAGUAAA[dU][dU]3′) (76) were synthetized by Sigma.

### Virus production

Viral particles were produced by Turbofect (Invitrogen) transfection of 293T cells following manufacturer instructions. Forty-eight hours after transfection, supernatants containing the viral particles were collected and filtered with a 0.45-μm filter to remove cell debris. The amounts of viruses produced was quantified by Gag-p24 ELISA.

### Infection experiments

Seventy-two hours after siRNA transfection, HeLa cells were infected with the HIV-1 strain NL4-3 (NIH AIDS Research and Reference Reagent Program, Division of AIDS, NIAID) pseudotyped with the vesicular stomatitis virus G glycoprotein (NL4-3-VSVG). Sixteen hours after infection, the viral input was removed. Twenty-four hours after, both viruses released in the supernatant and infected cells were analyzed by Western-blot and Gag-p24 ELISA. For the kinetics of Gag-p24 release in the supernatant of infected HeLa cells, viral input was removed twenty-four hours post-infection, and fresh medium was added. From this time point onward, viral supernatants were collected at 4 – 6 – 8 and 24 hours to perform Gag-p24 measurements using ELISA.

Monocyte-derived macrophages THP-1 cells were infected with HIV-1 strain NL4-3-VSVG. Sixteen hours after infection, viral input was removed. Forty-eight hours after, both viruses released in the supernatant and infected cells were analyzed by Western-blot and Gag-p24 ELISA. Jurkat T cells and activated primary CD4+ T cells were infected with the HIV-1 strain NL4-3-VSVG. Sixteen hours after infection, the viral input was removed. Twenty-four hours after, infected cells were analyzed by confocal spinning disk microscopy.

### Western Blot

Cells were lysed on ice for 30 min in Triton lysis buffer (50 mM Tris pH 8, 150 mM NaCl, 1 % Triton X-100 and protease inhibitors) and centrifuged at 10 000 g for 10 min. Infected cells and corresponding supernatants were lysed with 1 % Triton X-100 in PBS. Migration of proteins lysate was performed in 4-12 % gradient or in 10 % SDS-PAGE gels (Bio-Rad Laboratories), transferred onto PVDF membranes (Millipore), blocked and incubated with corresponding antibodies in 5 % milk in 50 mM Tris-HCl pH 8.0, 150 mM NaCl, 0.1 % Tween20, followed by horseradish peroxidase-coupled secondary antibodies (1:10,000, Jackson ImmunoResearch) and revealed by chemiluminescence (GE Healthcare).

### Gag-p24 ELISA

Infected cells and corresponding supernatants were inactivated with 1 % Triton X-100 in PBS. 96 well plates were washed with PBS and coated with an anti-Gag-p24 antibody (Nittobo, Cat#HIV-018–48304; 5 μg/mL) overnight at 37 °C. Plates were saturated with PBS 10 % FCS for 2 h at 37 °C. Samples were incubated for 2 h at 37 °C. An in-house-biotinylated detection anti-Gag-p24 antibody (clone 36/5.4A; Zeptometrix; 0.5 μg/mL) was added for 1 h at 37 °C followed by Streptavidin-conjugated Horse Raddish Peroxydase (HRP; BD Pharmingen; dilution 1:3000) for 1 h at 37 °C. O-phenylenediamine dihydrochloride (OPD; Thermo Fisher Scientific) solution was added for 30 min at room temperature in the dark and the reaction was stopped with H_2_SO_4_ 2 M. Optical density at 492 nm was measured. A standard curve was obtained using dilutions of a supernatant with known Gag-p24 content and used to infer absolute Gag-p24 concentrations. Each independent experiments were done in triplicate. Calculation of the HIV-1 release corresponds to the ratio of Gag-p24 in the supernatant to Gag-p24 in infected cells.

### Infectivity analysis

The infectivity of viral particles was assessed using P4C5 cells, which are HeLa CD4+ CCR5+ cells carrying an HIV-1 LTR–β-gal reporter cassette. One day prior to infection, 10^3^ cells/well were seeded in 96-well plates. Cells were then infected in triplicate with an amount of viruses corresponding to either 0.1 or 0.25 ng of Gag-p24. After 36 hours, cells were lysed in a solution containing PBS, 0.1 % NP-40, and 5 mM MgCl_2_, followed by incubation with the β-gal substrate CPRG (Roche). Subsequently, the absorbance at 570 nm was measured.

### Purification of viruses

Supernatants containing the viral particles were collected and filtered with a 0.45-μm filter to remove cell debris. The viral particles were further purified by ultracentrifugation of the filtered supernatants at 100,000 g for 1 hour. The resulting viral pellet was resuspended in PBS with agitation for 1 hour to ensure even resuspension. Both viral supernatants and ultracentrifuged viral particles were aliquoted and stored at -80°C until further use.

### Cell viability assay

Cell viability was quantified with the MTT assay (SIGMA, M2128). 100 µl of the cell suspension were pipetted into wells of a 96-well round-bottom plate, with triplicate samples prepared for each condition. Subsequently, 10 µl of MTT solution at a concentration of 7 mg/ml was added to each well. The plate was then incubated for 3 to 5 hours at 37°C in a 5% CO_2_ atmosphere. After incubation, 100 µl of acidic isopropanol (0.04 N) were added to each well to solubilize the formed crystals and inactivate viruses. Absorbance was measured at 540 nm.

### Transmission electron microscopy

Cells grown on coverslips were fixed in 80 mM PIPES, 5 mM EGTA, 2 mM MgCl_2_, 0.25 % triton, 2.5 % glutaraldehyde (Sigma Aldrich) and 1 % PFA for 20 min. Post fixation was done with 1 % osmium tetroxide (Merck) and 1.5 % ferrocyanide (Sigma Aldrich) in 0.1 M HEPES. After dehydration by a graded series of ethanol, cells on coverslips were infiltrated with epoxy resin. After polymerization thin sectioning was done using a Leica UC 7 microtome (Leica microsystems). The 70 nm sections were collected on formvar coated slot grids (EMS) and were contrasted with 4 % uranyl acetate and Reynolds lead citrate. Stained sections were observed with a Tecnai spirit FEI operated at 120 kV. Images were acquired with FEI Eagle digital camera.

### Correlative light and scanning electron microscopy

HeLa cells transfected with a NL-4.3 Gag-GFP provirus (kind gift from B. Müller and H. G. Kräusslich) (77) were platted on glass bottom dishes with a gridded coverslip (MatTek) to select cells producing viruses using fluorescent light microscope and subsequently to localize the same cells in SEM. The method for SEM preparation is described in details in ref. (78). Briefly, infected cells were fixed for 1 h in PHEM (60 mM PIPES pH 6.9; 25 mM HEPES; 2 mM MgCl_2_; 10 mM EGTA) containing 2.5 % Glutaraldehyde and 1 % PFA and washed three times with PHEM. Cells were imaged using an inverted Eclipse Ti-E microscope (Nikon) to identify and locate infected cells and their location on the grid. Cells were then post-fixed with 2 % osmium tetroxide (OsO_4_) in 1 M HEPES buffer at room temperature for 1 h and washed with distilled water. The samples were dehydrated by transfers into successive baths of ethanol (10, 25, 50, 75 and 95 %, 5 min each, and two times 100 %, 10 min each) and dried into the critical point dryer’s chamber (Leica EM CDP300). Then, samples were coated with 8 nm of gold/palladium (high-resolution ion beam coater, Gatan Model 681). The images were acquired with a JEOL JSM 6700 F field emission scanning electron microscope.

### Immunofluorescence and image acquisition with spinning disk confocal microcopy

Cells were grown on coverslips and fixed with paraformaldehyde (PFA) 4 % for 20 min at room temperature. Cells were then permeabilized and blocked with PBS containing 0.2 % bovine serum albumin (BSA) and 0.1 % saponin, and successively incubated for 1 h at room temperature with primary (see Antibodies section and *SI Appendix*, Table S1) and secondary antibodies (1:500, Jackson Laboratory) diluted in PBS containing 0.2 % BSA and 0.1 % saponin before DAPI staining for 5min (0.5 mg/mL, Serva). For F-actin staining, Alexa Fluor 561 or 647 Phalloidin (1:2000, Invitrogen) were added with the secondary antibodies. Cells were mounted in Fluoromount G (Southern Biotech).

Images were acquired with an inverted Nikon Eclipse Ti-E microscope equipped with a CSU-X1 spinning disk confocal scanning unit (Yokogawa) coupled to a Prime 95S scientific complementary metal-oxide semiconductor (sCMOS) camera (Teledyne Photometrics).

For Jurkat and activated primary CD4+ T cells, images were acquired with the LIVE-SR super resolution module (GATACA-Systems) added on the CSU-X1 spinning disk.

### Immunofluorescence and image acquisition with 3D STORM (STochastic Optical Reconstruction Microscopy)

Infected HeLa cells were grown on 22*22 mm 1.5H coverslips (EMS) and fixed in 80 mM PIPES, 5 mM EGTA, 2 mM MgCl_2_, 0.25 % triton, 0.5 % glutaraldehyde and 1 % PFA for 20 min. Glutaraldehyde was quenched with 0.1 % NaBH_4_ in PBS for 7 min. Cells were then permeabilized and saturated with PBS containing 0.22 % gelatin, 0.1 % Triton X-100 for 1h. Cells were incubated for 1 h at room temperature with anti-Gag antibody, then overnight with secondary antibodies (CF568) and Alexa Fluor 647 Phalloidin (ThermoFisher Scientific) diluted in PBS. Coverslips were incubated with a switching buffer prepared freshly before imaging (Tris 50 mM, NaCl 10 mM, 10 % glucose, Glucose oxidase 0.5 mg/mL, Catalase 50 µg/mL, MEA 50mM pH 8.3).

The super-resolved images were acquired on an Elyra 7 3D SMLM microscope (Carl Zeiss, Germany) in HILO illumination mode using a 63X/1.4 oil objective with a 1.518 refractive index oil (Carl Zeiss) and two sCMOS PCO Edge 4.2 cameras for the detection. The cameras were aligned using an internal pattern present in the optical path and chromatic aberrations were corrected for each image using 100 nm tetraspeck beads added to the cells. 3D STORM was performed using ‘double phase ramp’ optical component and calibrated with fluorescent beads. All processing was performed with the Zen software. The localization of individual molecules was determined using a peak mask size of 17 pixel, a peak intensity to noise of 5 and overlapping molecules were counted by setting a maximum cluster size of 10 pixels. Drift correction was corrected using model-based algorithm. Final images were generated with a resolution of 50 nm in Z and 10 nm X and Y.

### Image and data analysis

Images were mounted with the ImageJ software (NIH).

To quantify the number of budding regions per cell and the size of these regions (Fig. 2B, 5B, 6C and *SI Appendix*, Fig. S2B), images were processed using the “spot detector” plugin from the image analysis ICY software (Institut Pasteur, France-BioImaging) (79).

To quantify F-actin levels in Gag-positive and Gag-negative regions (Fig. 4B), orthogonal views of Z-stack images were analyzed using ImageJ. Mean intensity values were extracted from manually drawn ROIs for each channel of interest. The scan lines were obtained using an ROI (straight line), with a width corresponding to the actin cortex. Intensity values were normalized to Gag-negative regions.

To quantify the variations of F-actin intensity in Gag-positive regions compared to adjacent Gag-negative regions (Fig. 4C), orthogonal views of Z-stack images were analyzed using ImageJ. The scan lines were obtained using an ROI (straight line), with a width corresponding to the actin cortex. Variation of intensity of each channel of interest were analyzed using the Plot Profile function from ImageJ.

Mander’s overlap coefficient (ImageJ software) was used to quantify the colocalization between Gag and CHMP4B (*SI Appendix*, Fig. S2C).

Schematic illustrations were created with BioRender.com.

### TIRF microscopy

Live-cell TIRF microscopy in Figure 3C was carried out with an inverted Zeiss ELYRA PS.1 microscope (Carl Zeiss) using a αPlan-Apochromat x100/1.46 oil immersion objective and an EMCCD 512×512 Andor 897 camera driven by Zen black software. Cells were imaged 6–8 h after transfection of the plasmid encoding HIV-1 Gag-mCherry. The microscope was enclosed in a chamber and all imaging was carried out at 37 °C, 5 % CO_2_. Dual-color TIRF imaging of GFP and mCherry was achieved by exciting GFP with the 488nm laser (100 mW) and red fluorescent proteins with a 561nm laser (100 mW). Time-lapse movies were acquired over a 45 min period with one image every 30 s. Images were analyzed with the ImageJ software. Images were then converted into RGB 8-bit images using ImageJ software and processed with the Smooth function of ImageJ.

### Immunoprecipitation

For immunoprecipitations of endogenous proteins, HeLa cells were infected with NL4-3-VSVG and 24 hours after infection, viral input was removed and the cells were let for 24 hours. Cells were lysed in a buffer containing 50 mM Tris pH 7.4, 150 mM NaCl, 5 mM EGTA, and 1 % Triton X-100. Post-nuclear supernatants were incubated with Dynabeads Protein A (#10002D, Invitrogen) for 30 min (preclarification). Supernatants were then incubated with Dynabeads Protein A loaded with either anti-MICAL1 antibody (Proteintech, #14818-1-AP, 2μg) or loaded with control rabbit IgG (Millipore, #12-370, 2μg) for 1 h at 4 °C. After three washes in lysis buffer, proteins were resuspended in Laemmli buffer and boiled for 10 min at 95 °C.

For GFP immunoprecipitations, HeLa cells were transfected with either GFP or NL-4.3 Gag-GFP provirus and cells were lysed in a buffer containing 20 mM Tris pH 7.4, 150 mM NaCl, 5 mM EDTA and 1 % Triton X-100 36 hours post-transfection. Post-nuclear supernatants were incubated with GFP-Trap Magnetic Agarose beads (Chromotek, gtma-20) for 1 h at 4°C. After four washes in lysis buffer, proteins were resuspended in Laemmli buffer and boiled for 10 min at 95 °C.

### Yeast two-hybrid experiments

Yeast two-hybrid experiments were performed by co-transforming *the Saccharomyces cerevisiae* reporter strain L40 with either pGAD-empty, pGAD-MICAL1 (MICAL1 full length), pGAD-MICAL1^Cter^ (MICAL1^879–1067^), pGAD-MICAL1^Nter^ (MICAL1^1–843^) or pGAD-TSG101 together with either pLex empty, pLex-Rab35^Q67L^ or pLex-Gag HIV-1. Transformed yeast colonies were selected on DOB agarose plates without Tryptophane and Leucine. Colonies were picked and grown for 3 days on DOB agar plates with Histidine to select co-transformants and without Histidine to detect interactions.

### In vitro single actin filament assays

#### Proteins

Skeletal muscle actin (Uniprot P68135) was purified from rabbit muscle acetone powder following the original protocol from ref. (80), as described in ref. (81). Spectrin-actin seeds from human red blood cells were purified as described in ref. (81). Arp2/3 complex was purified from sheep brains as described in ref. (82). Recombinant N-terminal GST tag human N-WASP-VCA was purified as described in ref. (83).Recombinant monooxygenase catalytic domain of MICAL1 (MICAL1^1-499^) was purified as described in ref. (34). MICAL1^1-499^ was always used with its cofactor nicotinamide adenine dinucleotide phosphate (NADPH, 60 µM). HIV-1 Gag (NL4.3) recombinant protein has been purified from E. coli following the protocol described in (84).

#### Protein labeling

Actin was labeled on the surface lysine 328, using Alexa Fluor 568 (Thermo Fisher) or ATTO 488 NHS ester (Atto-tec) as described in ref. (85).

#### Buffers

All microfluidics experiments were carried out in F-buffer composed of 5 mM Tris HCl pH 7.0, 50 mM KCl, 1 mM MgCl_2_, 0.2 mM EGTA, 0.2 mM ATP, 10 mM DTT, 1 mM DABCO, 0.1 % Bovine Serum Albumin (BSA) at 25°C. In open-chamber experiments, the F-buffer was supplemented with 0.2 % Methyl cellulose.

#### Data acquisition

Observations were carried out on a Nikon TiE inverted microscope equipped with a 60x oil-immersion objective and an Evolve 512 EMCCD camera (Photometrics). A total internal reflection fluorescence (TIRF) illumination setup (iLAS2, Gataca-system), with 488 and 561 nm tunable lasers (maximum power of 100 mW each) was used. The laser exposure time for both channels was 200 ms. The manipulation and adjustment of the TIRF setup were done using Metamorph software (Version 7.10.4.407). Image analysis was performed using Fiji software.

#### Open chambers experiments

Experiments in open chambers (Fig. 5C), made from clean coverslips assembled with double-sided tape, were carried out as follows. First, the microchamber was passivated using BSA 5 % for 10 min and rinsed with F-buffer. Then, a mix of pre-polymerized mother filaments (formed from 1 μM 10 % AlexaFluor-568-labeled G-actin incubated in F-buffer in an Eppendorf tube for 2 hours, then diluted 40-fold in F-buffer) and branching solution (20 nM Arp2/3 complex, 50 nM VCA, 0.4 μM 10 % ATTO-488-labeled G-actin) was injected into the chamber. This solution was left in the chamber for 4 min, to allow the nucleation of branches. The solution was then replaced by a solution containing 0.6 µM 10 % AlexaFluor-568 labeled G-actin, in the presence or absence of 50 nM MICAL1^1-499^ and 60 µM NADPH. The branched filaments that remained in the chamber, close to the coverslip, were then monitored over time at a rate of 1 image every 5 sec.

#### Microfluidics experiments

Experiments in microfluidics chambers (Fig. 5D and 5E) were carried out using Poly-Dimethyl-Siloxane (PDMS, Sylgard) microchambers and a microfluidics device (MFCS and FLOW UNIT, Fluigent), based on the initial protocol from ref. (47) described in ref. (86). Spectrin-actin seeds were injected in the microfluidics chamber at 20 pM for 2 minutes. Then the surface was rinsed and passivated using 5 % BSA for 10 minutes. Next, mother filaments were polymerized by flowing in 0.6 μM 10 % ATTO488–labeled G-actin for 5 to 10 min. Mother filaments were exposed to the branching solution (20 nM Arp2/3 complex, 50 nM VCA, 0.4 μM 10 % AlexaFluor-568 labeled G-actin) to nucleate branches. Next mother filaments and branches were exposed to a solution containing 0.6 µM 10 % AlexaFluor-568 labeled G-actin, in the presence or absence of MICAL1^1-499^ (50 nM or 10 nM) and NADPH (60 µM). Images were acquired at a rate of 1 frame every 5 sec. The dissociation of filament branches and branch renucleation events were counted over time. Since purified Gag is stored at pH 8 with 1M NaCl, we modified our F-buffer to pH 7.4, supplemented with 200 mM NaCl for in vitro experiments monitoring the effect of Gag on debranching (*SI Appendix*, Fig. S4E). We first diluted Gag to 4 µM in its storage buffer, and then switched it to modified F-buffer in two dialysis steps using MINI Dialysis Devices (Thermo Scientific). After each dialysis step, we checked by ultracentrifugation and light absorption spectrometry (DeNovix) that no detectable aggregates were formed. Branch dissociation experiments were performed in microfluidics chambers as described above, except for the following differences: mother filaments were polymerized with 10% Alexa488-labeled G-actin; branch dissociation and renucleation, in the presence or absence of Gag, were monitored in modified F-buffer; images were acquired every 10 seconds.

#### Calculation of the force experienced by branches

The tension exerted on branches is a result of the friction of the fluid flow applied to the filaments. This tensile force was previously calibrated by ref. (87) and is proportional to the length of the branch, the flow velocity along the branch, and the friction force coefficient of the solution on the actin filament (η^actin^ = 6. 10-4 pN.µm^-2^.s). The initial average force and the force increase rate were calculated for each population of branches.

### Statistics

All plots and statistical tests were performed using the GraphPad Prism software. The presented values are displayed as mean ± SD from at least 3 independent experiments and the test used is indicated in the figure legends. In all statistical tests, p-value > 0.05 was considered as non-significant. P-values are indicated in the figures.

## DATA AVAILABILITY STATEMENT

All data associated with this paper are included in the manuscript and *SI Appendix*.

## AUTHOR CONTRIBUTIONS

T.S., N.C. and S.F conceived, carried out and analyzed the experiments presented in Figs. 1, 2, 3, 4, 5A, 5B, 6 and *SI Appendix*, Figs. S1, S2, S3, S4A, S4B, S4C and S5; F.G., H.W., A.J. and G.R-L. conceived, carried out and analyzed the experiments presented in Fig. 5C, 5D, 5E and *SI Appendix*, Fig. S4D and S4E; L.W., S.B. and M.R produced purified Gag; F.C. carried out and analyzed the experiments presented in *SI Appendix*, Fig. S4F. A.S. carried out and analyzed the experiments presented in Fig. 4D and 4E; M. M-N carried out the experiments presented in Fig. 3A and 3B; S.F., N.C., O.S., G.R-L, A.E. secured funding and supervised the work; S.F., N.C. and A.E. wrote the manuscript with the help of O.S, G. R-L, T.S.

## ACKNOWLEDGEMENTS

We thank P. Benaroch, C. Berlioz-Torrent, R. Dibsy and T. Wai for critical reading of the manuscript; the Echard Lab members for helpful discussions; the Romet/Jegou Lab members for help with in vitro experiments, P. Benaroch for plasmids. This work has been supported by Institut Pasteur, CNRS, ANR RedoxActin and Fondation pour le Recherche Médicale (EQU202103012627) to A.E., ANRS-21020 AP2020-2 to S.F.; Institut Pasteur, Urgence COVID-19 Fundraising Campaign of Institut Pasteur, Fondation pour la Recherche Médicale (FRM), ANRS, the Vaccine Research Institute (ANR-10-LABX-77), Labex IBEID (ANR-10-LABX-62-IBEID), ANR / FRM Flash Covid PROTEO-SARS-CoV-2, ANR Coronamito, HERA european funding (Durable consortium), LEAPS funding, ANRS-21020 AP2020-2 to N.C. and O.S.; ANR Redoxactin and Fondation pour la Recherche Medicale (EQU202203014630) to G.R-L; T.S. received a PhD fellowship from SIDACTION 2021-1-FJC-13007. We acknowledge the Ecole Doctorale Frontières de l’Innovation en Recherche et Education—Programme Bettencourt to T.S; F.G. received a post-doctoral fellowship from FRM (EQU202203014630). UTechS PBI and UTechS UBI are part of the France–BioImaging infrastructure network (FBI) supported by the French National Research Agency (ANR-10-INBS-04; Investments for the Future), and acknowledges support from Institut Pasteur, ANR/FBI, the Région Ile-de-France (program ‘Domaine d’Intérêt Majeur-Malinf’ and DIM1HEALTH) and the French Government Investissement d’Avenir Programme— Laboratoire d’Excellence ‘Integrative Biology of Emerging Infectious Diseases’ (ANR-10-LABX-62-IBEID) for the use of ELYRA PS1 LSM780 and ELYRA7 microscopes.

---

**Figure S1:**
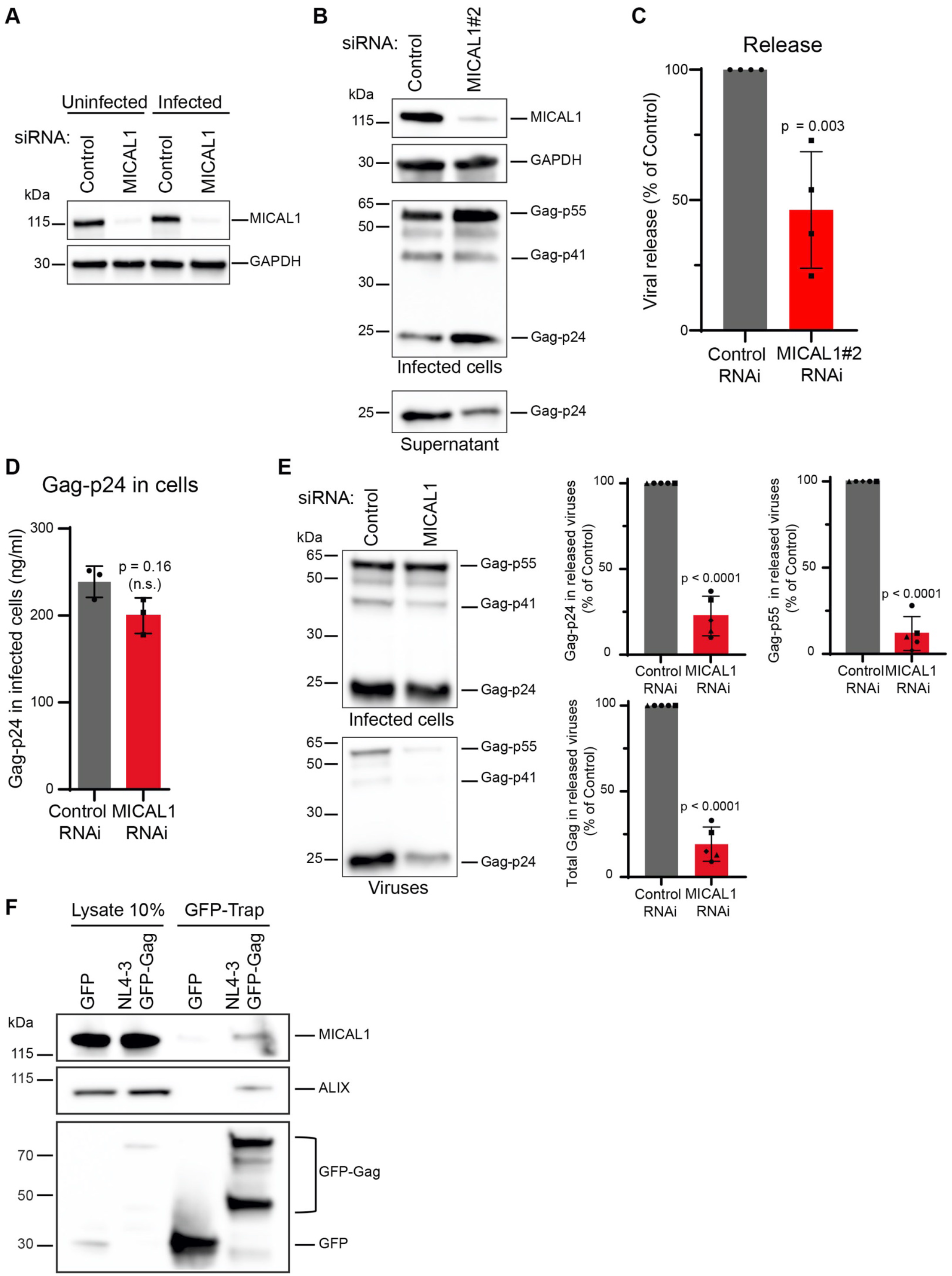
Viral release defects upon MICAL1 depletion. **(A)** Western blot analysis of MICAL1 in either uninfected or infected HeLa cells treated with indicated siRNAs. Loading control: GAPDH. This experiment was repeated at least 3 times independently with similar results. **(B)** Western blot analysis of HIV-1 Gag products in infected HeLa cells treated with indicated siRNAs (panels Infected cells) and supernatant (panel Supernatant). Loading control: GAPDH. Equal volumes of cell and supernatant samples were loaded. This experiment was repeated at least 3 times independently with similar results. **(C)** Quantification by Gag-p24 ELISA of HIV-1 release in HeLa cells treated with either Control or MICAL1#2 siRNAs. Results were normalized to control RNAi conditions (set at 100 %). Error bars represent SD calculated from 4 independent experiments, each done in triplicate. Two-tailed unpaired Student’s t-test. **(D)** Quantification by ELISA of Gag-p24 in infected HeLa cells treated with either Control or MICAL1 siRNAs (see corresponding Gag-p24 quantification in the supernatant in Fig. 1C). Error bars represent SD calculated from one experiment done in triplicate. Two-tailed unpaired Student’s t-test. **(E)** Left panels: Western blot analysis of HIV-1 Gag products in infected HeLa cells treated with indicated siRNAs (panel Infected cells) and corresponding purified viruses (panel Viruses). Equal volumes of cell and virus samples were loaded. This experiment was repeated 5 times with similar results. Right panels: Error bars represent SD calculated from 5 independent experiments. Two-tailed unpaired Student’s t-test. **(F)** HeLa cells were transfected with either GFP (control) or NL4-3-Gag-GFP provirus, and GFP was immunoprecipitated and revealed with anti-GFP antibodies (bottom panel). Co-immunoprecipitated endogenous MICAL1 (top panel) and ALIX (middle panel) were revealed with anti-MICAL1 and anti-ALIX antibodies, respectively.

**Figure S2:**
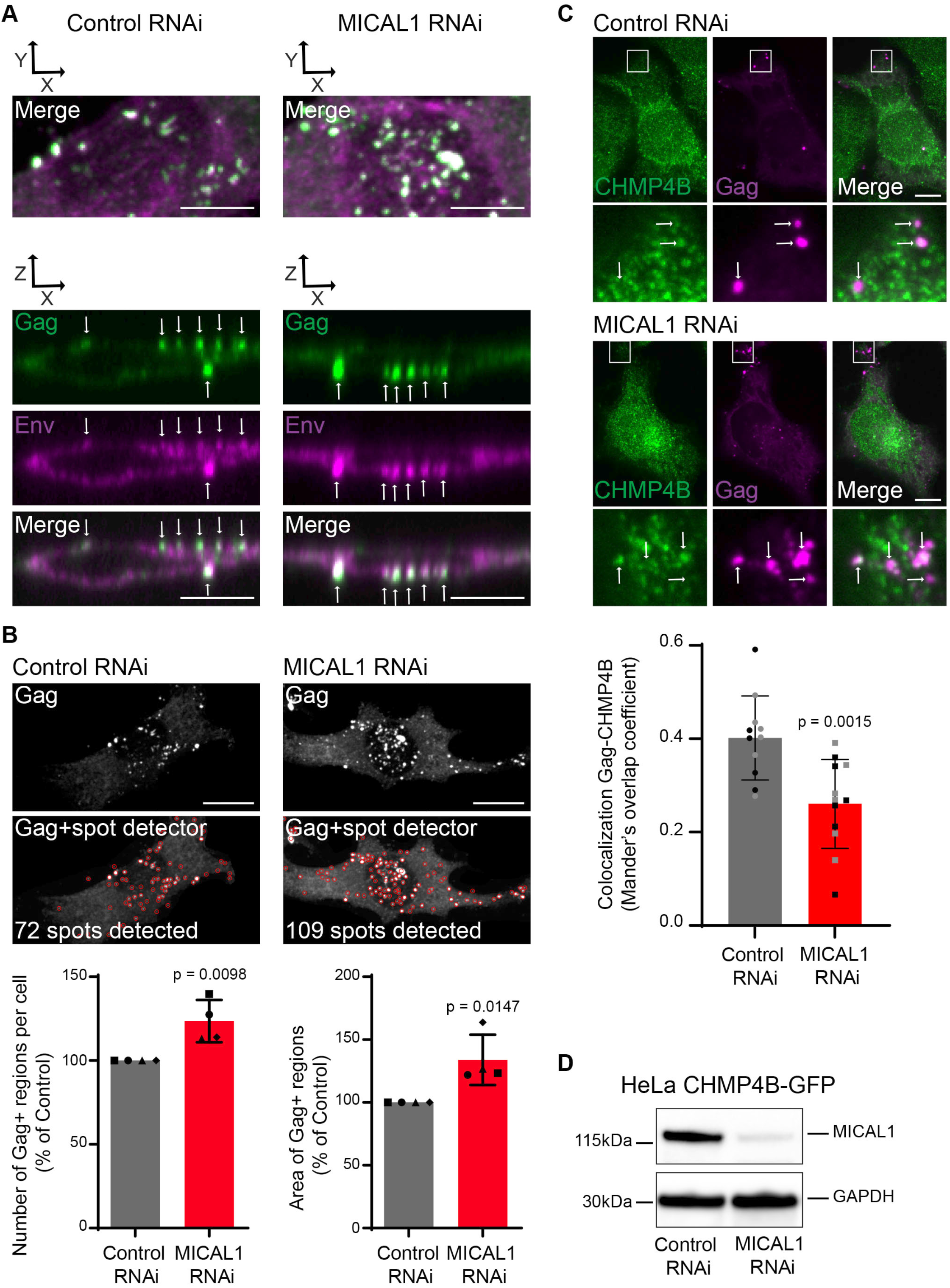
MICAL1 depletion induces an accumulation of HIV-1 budding particles at the plasma membrane and affects CHMP4B localization at budding regions. **(A)** Top panels: same images as in Fig. 2A (merged zoom, x,y views) of infected HeLa cells treated with indicated siRNAs and labeled with anti-Gag (green) and anti-Env (magenta) antibodies. Bottom panels: Corresponding Z stack images (x,z views) showing colocalization (arrows) of Gag and Env at the plasma membrane. Scale bars, 10 μm. **(B)** Top panels: results of the automatic spot detector analysis of the images presented in Fig. 2A. The number of detected spots corresponding to Gag+ regions are indicated. Bottom panels: Quantification of the number of Gag+ regions per cell and of the area of individual Gag+ regions automatically detected by spot detector in Control and MICAL1-depleted cells. n = 145 cells (Control RNAi) and n = 154 cells (MICAL1 RNAi). Error bars represent SD calculated from 4 independent experiments. Two-tailed unpaired Student’s t-test. **(C)** Top panels: Spinning disk confocal images of infected HeLa cells treated with indicated siRNAs and labeled with anti-CHMP4B antibody (green) and anti-Gag antibody (magenta). Arrows indicate Gag+ regions. Scale bars, 10 μm. Bottom panel: quantification of Gag-CHMP4B colocalization using Mander’s overlap coefficient (Error bars represent SD, 11 cells per condition, 2 independent experiments). Two-tailed unpaired Student’s t-test. **(D)** Western blot analysis of endogenous MICAL1 in a HeLa cell line that stably expresses CHMP4B-GFP. Loading control: GAPDH.

**Figure S3:**
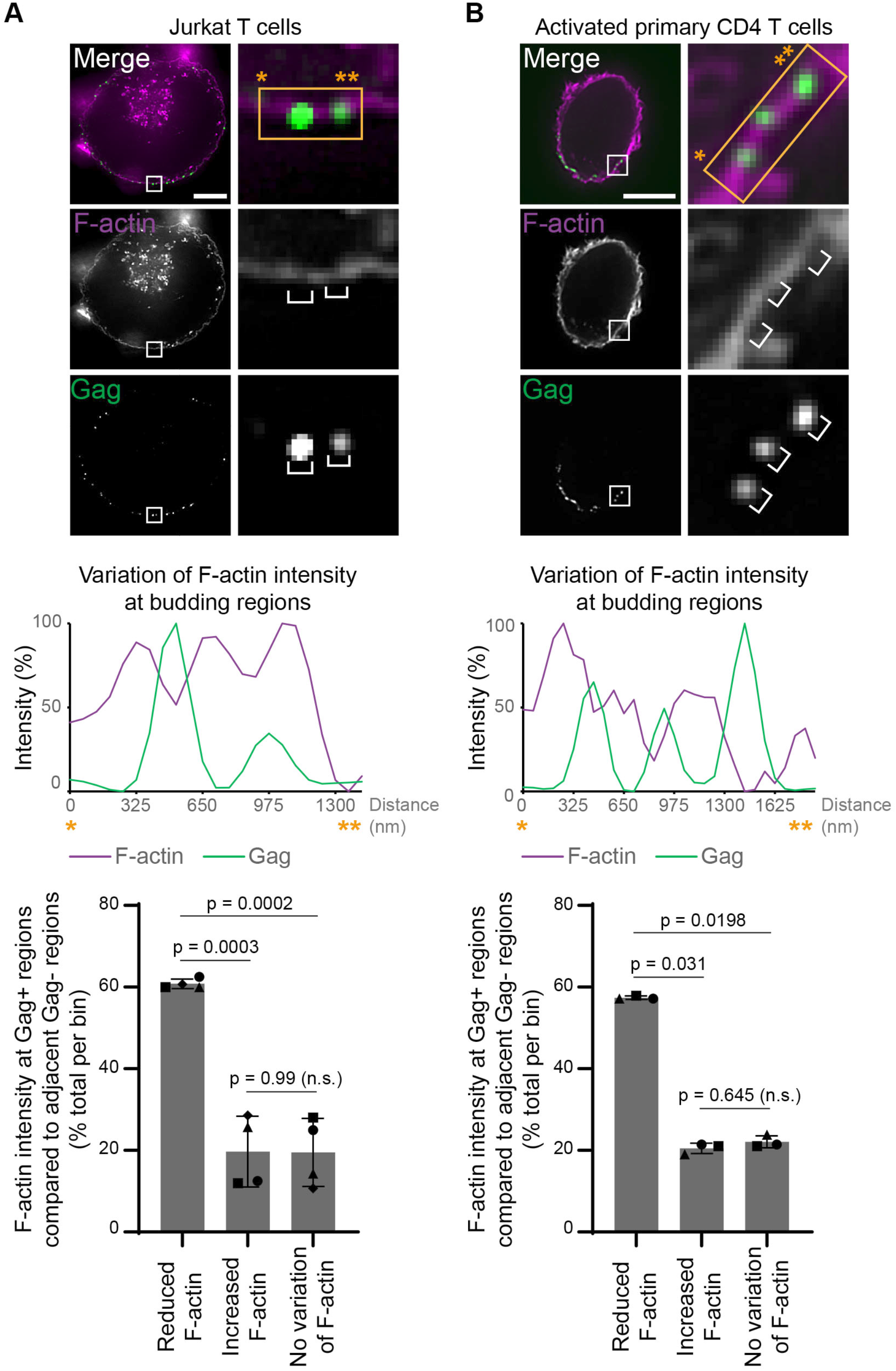
HIV-1 budding preferentially occurs in areas of the plasma membrane with low cortical F-actin levels in T lymphocytes. **(A)** Top panels: Spinning disk confocal images of infected Jurkat T cells labeled with anti-Gag antibody (green) and fluorescent phalloidin (magenta). Middle panel; Plot profile of the Gag and Phalloidin intensity on corresponding zoom image. Bottom panel: quantification of F-actin intensity at Gag+ regions compared to adjacent Gag-regions (Error bars represent SD, 120 budding regions from 4 cells). One-way ANOVA multiple comparisons. **(B)** Same as in (A) for activated primary CD4+ T cell (Error bars represent SD, 54 budding regions from 3 cells). Scale bars, 5 μm.

**Figure S4:**
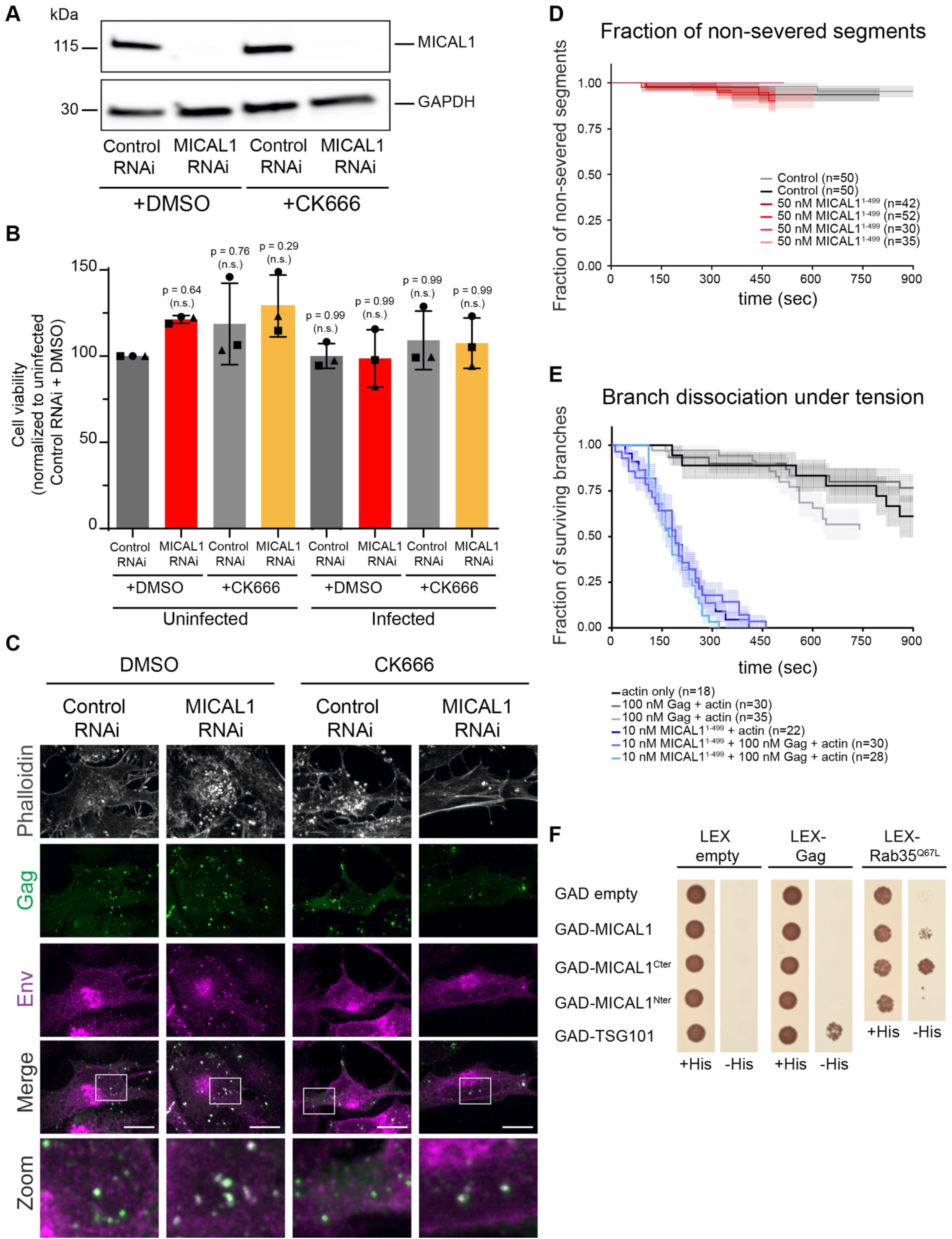
Effect of CK666 on HIV-1 budding regions in MICAL1-depleted cells and effect of Gag on MICAL1 debranching activity. **(A)** Western blot analysis of MICAL1 in HeLa cells treated with indicated siRNAs and either DMSO or 75 μM of CK666. Loading control: GAPDH. This experiment was repeated at least 3 times independently with similar results. **(B)** Cell viability of HeLa cells treated with indicated siRNAs and either DMSO or 75 μM CK666, both upon viral infection and in uninfected conditions. Error bars represent SD calculated from 3 independent experiments, each done in triplicate. One-way ANOVA multiple comparisons. **(C)** Z projection of spinning disk confocal images of infected HeLa cells treated with indicated siRNAs and either DMSO or CK666, and labeled with anti-Gag (green), anti-Env (magenta) antibodies and fluorescent phalloidin (grey). Scale bars, 10 μm. **(D)** Using the same filament populations as in Fig. 5C, the severing events occurring over the first 6 pixels of the filament branches were monitored over time. These 6-pixel segments (1.6 µm) correspond to the branch regions that were already polymerized when the debranching solution (without or with 50 nM MICAL1^1-499^ and 60 µM NADPH) was flowed into an open microchamber. Severing events within these segments were counted over time, and branch dissociation events were treated as censoring events using the Kaplan-Meier method. The shaded areas show the 65% confidence intervals. **(E)** Fraction of remaining (undissociated) actin filament branches as a function of time, in the presence or absence of 10 nM MICAL^1-499^ (plus NADPH), and with or without 100 nM Gag. The shaded areas show 65% confidence intervals. The renucleation ratios corresponding to branch populations under different conditions from top (actin only) to bottom (10 nM FAD + 100 nM Gag + actin) are 50 % (±12.5%), 40% (±9.7%), 53.3% (±12.8%), 40.9% (±10.4%), 36.6% (±8.7%), and 46.4% (±9.4%), respectively. **(F)** Yeast two hybrid assay using *S. cerevisiae* L40 reporter strain expressing indicated GAD and Lex fusion proteins, and grown on DOB selective medium with (+His) or without Histidine (-His). Absence of colony growth in DOB -Histidine indicates a lack of detection of interaction between the tested proteins.

**Figure S5:**
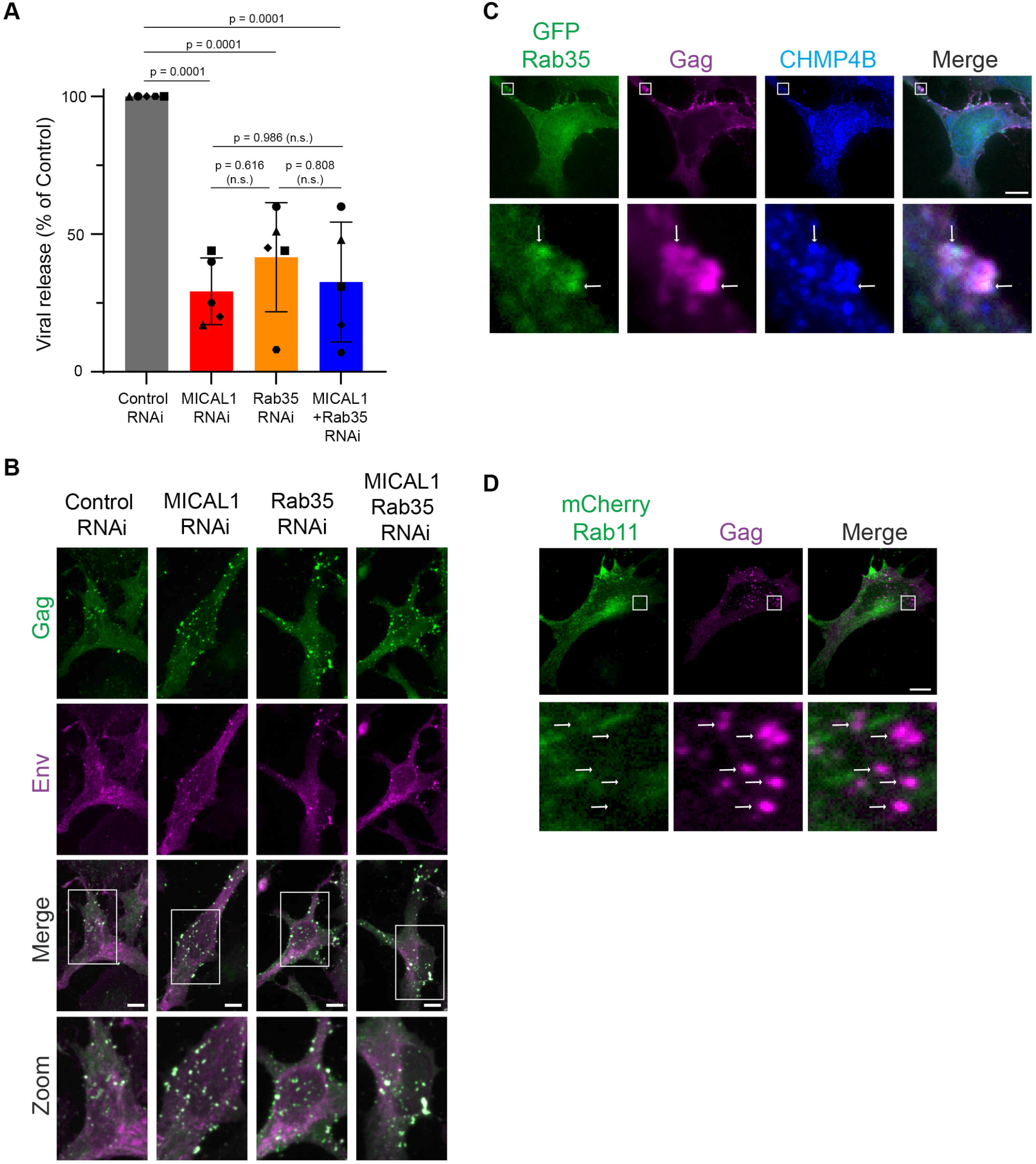
MICAL1 and its activator the GTPase Rab35 act in the same pathway in HIV-1 release. **(A)** Quantification of HIV-1 release by WB using anti-Gag antibodies of infected HeLa cells treated with indicated siRNAs. Error bars represent SD calculated from 5 independent experiments. One-way ANOVA multiple comparisons. **(B)** Z projection of spinning disk confocal images of infected HeLa cells treated with indicated siRNAs and labeled with anti-Gag (green) and anti-Env (magenta) antibodies. Scale bars, 10 μm. **(C)** Spinning disk confocal images of infected HeLa cells that express endogenous Rab35 tagged with GFP and stained with anti-Gag antibodies (magenta), GFP-Booster nanobodies (green) and anti-CHMP4B antibodies (blue). White arrows indicate Gag+ regions. Scale bars, 10 μm. **(D)** Spinning disk confocal images of infected HeLa cells transfected with plasmid encoding mCherry-Rab11 (green) and labeled with anti-Gag (magenta) antibodies. White arrows indicate Gag+ regions. Scale bars, 10 μm.

**Table S1:** Antibodies, dilutions and fixations used in this study.

| target protein | company | host | name/clone | dilution WB | dilution IF | fixation IF |
| --- | --- | --- | --- | --- | --- | --- |
| MICAL1 | Proteintech | rabbit | 14818-1-AP | 1/500 |  |  |
| Rab35 | Proteintech | rabbit | 11329-2-AP | 1/500 |  |  |
| GAPDH | Proteintech | mouse | 60004-1-Ig | 1/8000 |  |  |
| Gag | NIH | rabbit | ARP-4250 | 1/5000 | 1/2000 | PFA |
| Gag | NIH | mouse | ARP-3537 | 1/5000 |  |  |
| GFP | Proteintech | mouse | 66002-1-Ig | 1/2000 |  |  |
| Env | Institut Pasteur | mouse | Clone 110H |  | 1/500 | PFA |
| CHMP4B | Proteintech | rabbit | 13683-1-AP |  | 1/200 | MetOH |
| ALIX | Santa Cruz Biotechnology | mouse | sc-271975 | 1/1000 |  |  |

PFA: paraformaldehyde fixation
MetOH: methanol

## REFERENCES

1. D. G. Demirov, J. M. Orenstein, E. O. Freed, The late domain of human immunodeficiency virus type 1 p6 promotes virus release in a cell type-dependent manner. J Virol 76, 105–117 (2002).

2. J. E. Garrus et al., Tsg101 and the vacuolar protein sorting pathway are essential for HIV-1 budding. Cell 107, 55–65 (2001).

3. J. Martin-Serrano, T. Zang, P. D. Bieniasz, HIV-1 and Ebola virus encode small peptide motifs that recruit Tsg101 to sites of particle assembly to facilitate egress. Nat Med 7, 1313–1319 (2001).

4. L. VerPlank et al., Tsg101, a homologue of ubiquitin-conjugating (E2) enzymes, binds the L domain in HIV type 1 Pr55(Gag). Proc Natl Acad Sci U S A 98, 7724–7729 (2001).

5. U. K. von Schwedler et al., The protein network of HIV budding. Cell 114, 701–713 (2003).

6. J. G. Carlton, J. Martin-Serrano, Parallels between cytokinesis and retroviral budding: a role for the ESCRT machinery. Science 316, 1908–1912 (2007).

7. E. Morita et al., Human ESCRT and ALIX proteins interact with proteins of the midbody and function in cytokinesis. Embo J 26, 4215–4227 (2007).

8. M. Vietri, M. Radulovic, H. Stenmark, The many functions of ESCRTs. Nat Rev Mol Cell Biol 21, 25–42 (2020).

9. J. McCullough, A. Frost, W. I. Sundquist, Structures, Functions, and Dynamics of ESCRT-III/Vps4 Membrane Remodeling and Fission Complexes. Annu Rev Cell Dev Biol 34, 85–109 (2018).

10. E. J. Scourfield, J. Martin-Serrano, Growing functions of the ESCRT machinery in cell biology and viral replication. Biochem Soc Trans 45, 613–634 (2017).

11. G. Lerner, N. Weaver, B. Anokhin, P. Spearman, Advances in HIV-1 Assembly. Viruses 14 (2022).

12. W. I. Sundquist, H. G. Krausslich, HIV-1 assembly, budding, and maturation. Cold Spring Harb Perspect Med 2, a006924 (2012).

13. E. O. Freed, HIV-1 assembly, release and maturation. Nat Rev Microbiol 13, 484–496 (2015).

14. P. Chugh, E. K. Paluch, The actin cortex at a glance. J Cell Sci 131 (2018).

15. M. Kelkar, P. Bohec, G. Charras, Mechanics of the cellular actin cortex: From signalling to shape change. Curr Opin Cell Biol 66, 69–78 (2020).

16. A. M. Gautreau, F. E. Fregoso, G. Simanov, R. Dominguez, Nucleation, stabilization, and disassembly of branched actin networks. Trends Cell Biol 32, 421–432 (2022).

17. R. S. Kadzik, K. E. Homa, D. R. Kovar, F-Actin Cytoskeleton Network Self-Organization Through Competition and Cooperation. Annu Rev Cell Dev Biol 36, 35–60 (2020).

18. C. Jolly, I. Mitar, Q. J. Sattentau, Requirement for an intact T-cell actin and tubulin cytoskeleton for efficient assembly and spread of human immunodeficiency virus type 1. J Virol 81, 5547–5560 (2007).

19. G. Audoly, M. R. Popoff, P. Gluschankof, Involvement of a small GTP binding protein in HIV-1 release. Retrovirology 2, 48 (2005).

20. M. Gladnikoff, E. Shimoni, N. S. Gov, I. Rousso, Retroviral assembly and budding occur through an actin-driven mechanism. Biophys J 97, 2419–2428 (2009).

21. L. A. Carlson et al., Cryo electron tomography of native HIV-1 budding sites. PLoS Pathog 6, e1001173 (2010).

22. R. Dibsy, E. Bremaud, J. Mak, C. Favard, D. Muriaux, HIV-1 diverts cortical actin for particle assembly and release. Nat Commun 14, 6945 (2023).

23. S. A. Rahman et al., Investigating the role of F-actin in human immunodeficiency virus assembly by live-cell microscopy. J Virol 88, 7904–7914 (2014).

24. S. Stauffer et al., The nucleocapsid domain of Gag is dispensable for actin incorporation into HIV-1 and for association of viral budding sites with cortical F-actin. J Virol 88, 7893–7903 (2014).

25. T. Serrano, S. Fremont, A. Echard, Get in and get out: Remodeling of the cellular actin cytoskeleton upon HIV-1 infection. Biol Cell 10.1111/boc.202200085 (2023).

26. D. Dambournet et al., Rab35 GTPase and OCRL phosphatase remodel lipids and F-actin for successful cytokinesis. Nat Cell Biol 13, 981–988 (2011).

27. J. A. Schiel et al., FIP3-endosome-dependent formation of the secondary ingression mediates ESCRT-III recruitment during cytokinesis. Nat Cell Biol 14, 1068–1078 (2012).

28. S. Fremont, A. Echard, Membrane Traffic in the Late Steps of Cytokinesis. Curr Biol 28, R458–R470 (2018).

29. R. Kumar et al., DENND2B activates Rab35 at the intercellular bridge, regulating cytokinetic abscission and tetraploidy. Cell Rep 42, 112795 (2023).

30. N. V. G. Iannantuono, G. Emery, Rab11FIP1 maintains Rab35 at the intercellular bridge to promote actin removal and abscission. J Cell Sci 134 (2021).

31. S. J. Terry, F. Dona, P. Osenberg, J. G. Carlton, U. S. Eggert, Capping protein regulates actin dynamics during cytokinetic midbody maturation. Proc Natl Acad Sci U S A 115, 2138–2143 (2018).

32. J. Bai et al., Actin reduction by MsrB2 is a key component of the cytokinetic abscission checkpoint and prevents tetraploidy. Proc Natl Acad Sci U S A 117, 4169–4179 (2020).

33. C. Addi, J. Bai, A. Echard, Actin, microtubule, septin and ESCRT filament remodeling during late steps of cytokinesis. Curr Opin Cell Biol 50, 27–34 (2018).

34. S. Fremont et al., Oxidation of F-actin controls the terminal steps of cytokinesis. Nat Commun 8, 14528 (2017).

35. C. Rouyere, T. Serrano, S. Fremont, A. Echard, Oxidation and reduction of actin: Origin, impact in vitro and functional consequences in vivo. Eur J Cell Biol 101, 151249 (2022).

36. L. T. Alto, J. R. Terman, MICALs. Curr Biol 28, R538–R541 (2018).

37. S. S. Giridharan, S. Caplan, MICAL-family proteins: Complex regulators of the actin cytoskeleton. Antioxid Redox Signal 20, 2059–2073 (2014).

38. M. A. Vanoni, Structure-function studies of MICAL, the unusual multidomain flavoenzyme involved in actin cytoskeleton dynamics. Arch Biochem Biophys 632, 118–141 (2017).

39. S. Rajan, J. R. Terman, E. Reisler, MICAL-mediated oxidation of actin and its effects on cytoskeletal and cellular dynamics. Front Cell Dev Biol 11, 1124202 (2023).

40. G. Mercenne, S. L. Alam, J. Arii, M. S. Lalonde, W. I. Sundquist, Angiomotin functions in HIV-1 assembly and budding. Elife 4 (2015).

41. K. Inamdar et al., Full assembly of HIV-1 particles requires assistance of the membrane curvature factor IRSp53. Elife 10 (2021).

42. D. G. Demirov, A. Ono, J. M. Orenstein, E. O. Freed, Overexpression of the N-terminal domain of TSG101 inhibits HIV-1 budding by blocking late domain function. Proc Natl Acad Sci U S A 99, 955–960 (2002).

43. C. Addi et al., The Flemmingsome reveals an ESCRT-to-membrane coupling via ALIX/syntenin/syndecan-4 required for completion of cytokinesis. Nat Commun 11, 1941 (2020).

44. N. Jouvenet, P. D. Bieniasz, S. M. Simon, Imaging the biogenesis of individual HIV-1 virions in live cells. Nature 454, 236–240 (2008).

45. N. Jouvenet, M. Zhadina, P. D. Bieniasz, S. M. Simon, Dynamics of ESCRT protein recruitment during retroviral assembly. Nat Cell Biol 13, 394–401 (2011).

46. E. E. Grintsevich et al., Catastrophic disassembly of actin filaments via Mical-mediated oxidation. Nat Commun 8, 2183 (2017).

47. A. Jegou, et al., Individual actin filaments in a microfluidic flow reveal the mechanism of ATP hydrolysis and give insight into the properties of profilin. Plos Biol 9, e1001161 (2011).

48. F. Ghasemi et al., Regeneration of actin filament branches from the same Arp2/3 complex. Sci Adv 10, eadj7681 (2024).

49. N. G. Pandit et al., Force and phosphate release from Arp2/3 complex promote dissociation of actin filament branches. Proc Natl Acad Sci U S A 117, 13519–13528 (2020).

50. C. Cauvin et al., Rab35 GTPase Triggers Switch-like Recruitment of the Lowe Syndrome Lipid Phosphatase OCRL on Newborn Endosomes. Curr Biol 26, 120–128 (2016).

51. M. Fukuda, E. Kanno, K. Ishibashi, T. Itoh, Large scale screening for novel rab effectors reveals unexpected broad Rab binding specificity. Mol Cell Proteomics 7, 1031–1042 (2008).

52. A. G. Clark, O. Wartlick, G. Salbreux, E. K. Paluch, Stresses at the cell surface during animal cell morphogenesis. Curr Biol 24, R484–494 (2014).

53. M. Barrero-Villar et al., Moesin is required for HIV-1-induced CD4-CXCR4 interaction, F-actin redistribution, membrane fusion and viral infection in lymphocytes. J Cell Sci 122, 103–113 (2009).

54. N. H. Roy, M. Lambele, J. Chan, M. Symeonides, M. Thali, Ezrin is a component of the HIV-1 virological presynapse and contributes to the inhibition of cell-cell fusion. J Virol 88, 7645–7658 (2014).

55. H. Kamiyama et al., Role of Ezrin Phosphorylation in HIV-1 Replication. Front Microbiol 9, 1912 (2018).

56. H. Wu, H. G. Yesilyurt, J. Yoon, J. R. Terman, The MICALs are a Family of F-actin Dismantling Oxidoreductases Conserved from Drosophila to Humans. Sci Rep 8, 937 (2018).

57. R. J. Hung et al., Mical links semaphorins to F-actin disassembly. Nature 463, 823–827 (2010).

58. R. J. Hung, C. W. Pak, J. R. Terman, Direct redox regulation of F-actin assembly and disassembly by Mical. Science 334, 1710–1713 (2011).

59. S. Rajan et al., Disassembly of bundled F-actin and cellular remodeling via an interplay of Mical, cofilin, and F-actin crosslinkers. Proc Natl Acad Sci U S A 120, e2309955120 (2023).

60. C. Galloni et al., MICAL2 enhances branched actin network disassembly by oxidizing Arp3B-containing Arp2/3 complexes. J Cell Biol 220 (2021).

61. E. F. Schmidt, S. O. Shim, S. M. Strittmatter, Release of MICAL autoinhibition by semaphorin-plexin signaling promotes interaction with collapsin response mediator protein. J Neurosci 28, 2287–2297 (2008).

62. J. R. Terman, T. Mao, R. J. Pasterkamp, H. H. Yu, A. L. Kolodkin, MICALs, a family of conserved flavoprotein oxidoreductases, function in plexin-mediated axonal repulsion. Cell 109, 887–900 (2002).

63. S. S. Giridharan, J. L. Rohn, N. Naslavsky, S. Caplan, Differential regulation of actin microfilaments by human MICAL proteins. J Cell Sci 125, 614–624 (2012).

64. T. Vitali, E. Maffioli, G. Tedeschi, M. A. Vanoni, Properties and catalytic activities of MICAL1, the flavoenzyme involved in cytoskeleton dynamics, and modulation by its CH, LIM and C-terminal domains. Arch Biochem Biophys 593, 24–37 (2016).

65. A. Rai et al., bMERB domains are bivalent Rab8 family effectors evolved by gene duplication. eLife 5 (2016).

66. V. Chukkapalli, I. B. Hogue, V. Boyko, W. S. Hu, Ono, Interaction between the human immunodeficiency virus type 1 Gag matrix domain and phosphatidylinositol-(4,5)-bisphosphate is essential for efficient gag membrane binding. J Virol 82, 2405–2417 (2008).

67. V. Chukkapalli, A. Ono, Molecular determinants that regulate plasma membrane association of HIV-1 Gag. J Mol Biol 410, 512–524 (2011).

68. A. Ono, S. D. Ablan, S. J. Lockett, K. Nagashima, E. O. Freed, Phosphatidylinositol (4,5) bisphosphate regulates HIV-1 Gag targeting to the plasma membrane. Proc Natl Acad Sci U S A 101, 14889–14894 (2004).

69. C. Favard et al., HIV-1 Gag specifically restricts PI(4,5)P2 and cholesterol mobility in living cells creating a nanodomain platform for virus assembly. Sci Adv 5, eaaw8651 (2019).

70. P. Sengupta et al., A lipid-based partitioning mechanism for selective incorporation of proteins into membranes of HIV particles. Nat Cell Biol 21, 452–461 (2019).

71. W. D. Heo et al., PI(3,4,5)P3 and PI(4,5)P2 lipids target proteins with polybasic clusters to the plasma membrane. Science 314, 1458–1461 (2006).

72. K. Klinkert, A. Echard, Rab35 GTPase: a central regulator of phosphoinositides and F-actin in endocytic recycling and beyond. Traffic 10.1111/tra.12422 (2016).

73. K. Dicker, A. I. Jarvelin, M. Garcia-Moreno, A. Castello, The importance of virion-incorporated cellular RNA-Binding Proteins in viral particle assembly and infectivity. Semin Cell Dev Biol 111, 108–118 (2021).

74. K. Miyauchi, M. Marin, G. B. Melikyan, Visualization of retrovirus uptake and delivery into acidic endosomes. Biochem J 434, 559–569 (2011).

75. W. Deng et al., MICAL1 controls cell invasive phenotype via regulating oxidative stress in breast cancer cells. BMC Cancer 16, 489 (2016).

76. I. Kouranti, M. Sachse, N. Arouche, B. Goud, A. Echard, Rab35 regulates an endocytic recycling pathway essential for the terminal steps of cytokinesis. Curr Biol 16, 1719–1725 (2006).

77. B. Muller et al., Construction and characterization of a fluorescently labelled infectious human immunodeficiency virus type 1 derivative. J Virol 78, 10803–10813 (2004).

78. S. Fremont, A. Echard, Studying cytokinesis and midbody remnants using correlative light/scanning EM. Methods Cell Biol 137, 239–251 (2017).

79. F. de Chaumont et al., Icy: an open bioimage informatics platform for extended reproducible research. Nat Methods 9, 690–696 (2012).

80. J. A. Spudich, S. Watt, The regulation of rabbit skeletal muscle contraction. I. Biochemical studies of the interaction of the tropomyosin-troponin complex with actin and the proteolytic fragments of myosin. J Biol Chem 246, 4866–4871 (1971).

81. H. Wioland et al., ADF/Cofilin Accelerates Actin Dynamics by Severing Filaments and Promoting Their Depolymerization at Both Ends. Curr Biol 27, 1956–1967 e1957 (2017).

82. C. Le Clainche, M. F. Carlier, Actin-based motility assay. Curr Protoc Cell Biol Chapter 12, 12 17 11–12 17 20 (2004).

83. L. Cao, F. Ghasemi, M. Way, A. Jegou, G. Romet-Lemonne, Regulation of branched versus linear Arp2/3-generated actin filaments. EMBO J 42, e113008 (2023).

84. W. J. McKinstry et al., Expression and purification of soluble recombinant full length HIV-1 Pr55(Gag) protein in Escherichia coli. Protein Expr Purif 100, 10–18 (2014).

85. G. Romet-Lemonne, B. Guichard, A. Jegou, Using Microfluidics Single Filament Assay to Study Formin Control of Actin Assembly. Methods Mol Biol 1805, 75–92 (2018).

86. H. Wioland, F. Ghasemi, J. Chikireddy, G. Romet-Lemonne, A. Jegou, Using Microfluidics and Fluorescence Microscopy to Study the Assembly Dynamics of Single Actin Filaments and Bundles. J Vis Exp 10.3791/63891 (2022).

87. A. Jegou, T. Niedermayer, R. Lipowsky, M. F. Carlier, G. Romet-Lemonne, On phosphate release in actin filaments. Biophys J 104, 2778–2779 (2013).

